# mRNA Vaccine-Induced SARS-CoV-2 Spike-Specific IFN-γ and IL-2 T-cell Responses are Predictive of Serological Neutralization and are Transiently Enhanced by Pre-Existing Cross-Reactive Immunity

**DOI:** 10.1101/2024.05.28.596157

**Authors:** Philip Samaan, Chapin S. Korosec, Patrick Budylowski, Serena L. L. Chau, Adrian Pasculescu, Freda Qi, Melanie Delgado-Brand, Tulunay R. Tursun, Geneviève Mailhot, Roya Monica Dayam, Corey R. Arnold, Marc-André Langlois, Anjali Patel, Keelia Quinn de Launay, Jamie M. Boyd, Alyson Takaoka, Karen Colwill, Allison McGeer, Sharon Straus, Anne-Claude Gingras, Jane M. Heffernen, Mario Ostrowski

## Abstract

The contributions of SARS-CoV-2-specific T-cells to vaccine efficacy and durability are unclear. We investigated the relationships between mRNA vaccine-induced spike-specific IFN-γ and IL-2 T-cell responses, anti-spike/RBD IgG/IgA antibodies, and live virus neutralizing capacity in long-term-care-home staff doubly vaccinated with BNT162b2 or mRNA-1273. The impacts of pre-existing cross-reactive T-cell immunity to SARS-CoV-2 on cellular and humoral responses to vaccination were additionally assessed. Mathematical modelling of the kinetics of spike-specific IFN-γ and IL-2 T-cell responses over 6-months post-second dose was bifurcated into recipients who exhibited gradual increases (54% and 42%, respectively) with doubling times of 173 days, or decreases (46% and 58%, respectively) with half-lives of 115 days. Differences in kinetics did not correlate with any clinical phenotypes, although increases were proposed to be due to subclinical viral exposures. Serological anti-spike/RBD IgG/IgA antibody levels had otherwise decayed in all participants with half-lives of 55, 53, 76, and 59 days, respectively. Spike-specific T-cell responses induced at 2-6 weeks correlated with live viral neutralization at 6-months post-second dose, especially in hybrid immune individuals. Participants with pre-existing cross-reactive T-cell immunity to SARS-CoV-2 exhibited greater spike-specific T-cell responses, reduced anti-RBD IgA antibody levels, and a trending increase in neutralization at 2-6 weeks post-second dose. Non-spike-specific T-cells predominantly targeted SARS-CoV-2 non-structural protein at 6-months post-second dose in cross-reactive participants. mRNA vaccination was lastly shown to induce off-target T-cell responses against unrelated antigens. In summary, vaccine-induced spike-specific T-cell immunity appeared to influence serological neutralizing capacity, with only a modest effect induced by pre-existing cross-reactivity.

**IMPORTANCE:** Our findings provide important insights on the potential contributions of mRNA vaccine-induced spike-specific T-cell responses to the durability of neutralizing antibody levels in both uninfected and hybrid immune recipients. Our study additionally sheds light on the precise impacts of pre-existing cross-reactive T-cell immunity to SARS-CoV-2 on the magnitude and kinetics of cellular and humoral responses to vaccination. This study should ultimately inform the development of novel pan-coronavirus vaccines and vaccine regimens that can maximize the durability and breadth of protection against both current and future human coronaviruses of concern.

## INTRODUCTION

The COVID-19 pandemic, caused by the SARS-CoV-2 virus, continues to pose a serious threat to public health since its emergence in Wuhan, China in December 2019. Immunological correlates of protection against infection, symptomatic disease, and death are still unclear (1). Investigating these correlates can aid in the identification of surrogate markers that can be used to predict SARS-CoV-2 vaccine efficacy and durability on an individual basis.

T-cells are key players in the adaptive immune response to invading pathogens (2). In COVID-19, the earlier induction of robust SARS-CoV-2-specific T-cell responses during acute infection are well correlated with viral control and milder disease course (3–6). The importance of T-cells in viral clearance is likewise highlighted within the context of vaccination. As SARS-CoV-2 rapidly evolved to evade neutralizing humoral responses over time, vaccine regimens were rendered progressively less efficacious against symptomatic infection (7–9). Yet, the same vaccine regimens maintained a high degree of protection against severe disease (7–9), which aligned with accompanying reports indicating that vaccine-induced spike-specific T-cell responses across SARS-CoV-2 variants of concern (VOCs) are mostly preserved (10, 11). Accordingly, much interest has been garnered in the role of T-cells in vaccination against SARS-CoV-2.

CD4^+^ T-cells are critical for directing the adaptive immune response to viral pathogens. Firstly, CD4^+^ T-cells can provide indirect help to CD8^+^ T-cells by licencing antigen-presenting dendritic cells, which in turn become more efficient at activating CD8^+^ T-cells (12). Activated CD8^+^ T-cells sequentially proliferate and differentiate into cytotoxic T lymphocytes that kill virally infected target cells (12). A significant negative moderate correlation between SARS-CoV-2-specific CD8^+^ T-cell responses and peak disease severity in COVID-19 patients has notably been observed in a previous report (5). In this regard, memory CD4^+^ T-cells may be crucial for the preservation of protection against severe disease conferred by vaccine regimens across VOCs (9, 13, 14). Secondly, CD4^+^ T-cells provide essential help to B-cells for the development of neutralizing antibodies (15, 16). Previous studies have reported that the potency and earlier induction of SARS-CoV-2-specific neutralizing antibodies are predictive of milder disease course and survival in COVID-19 patients (17, 18). Accordingly, the conservation of SARS-CoV-2-specific T-cell responses across VOCs may have preserved the ability to quickly induce robust neutralizing humoral responses against infection (19), preventing severe disease.

Within the context of natural infection, several previous studies have already shown significant positive associations between SARS-CoV-2-specific CD4^+^ T-cell responses and serological antibody levels and neutralization (4, 5, 20–22). However, these associations have been evaluated in very few studies to date within the context of vaccination (23, 24). Most importantly, the precise relationships between the kinetics of vaccine-induced spike-specific CD4^+^ T-cell responses and neutralizing antibody levels have not been thoroughly investigated.

It was previously found that the neutralizing activity of acute and convalescent plasma/serum is mostly facilitated by spike- and RBD-specific IgM and IgG1 antibodies (25). Serological IgA antibodies were also reported to contribute to neutralizing activity (25) and have been observed to play a potential role in protection against infection (26). Considering the collective contributions of IFN-γ and IL-2 secreted by CD4^+^ T helper cells to B-cell proliferation, differentiation, and survival along with antibody class-switching to IgG and IgA (20, 27–32), we hypothesize that spike-specific IFN-γ and IL-2 T-cell responses to vaccination will enhance neutralizing antibody levels against SARS-CoV-2.

Previous exposure to human common-cold coronaviruses (HCoVs) may potentially impact both spike-specific CD4^+^ T-cell and neutralizing humoral responses to vaccination due to their significant homologies with SARS-CoV-2 (33). Multiple studies have shown that at the beginning of the COVID-19 pandemic, 20-50% of healthy, unexposed individuals exhibited pre-existing cross-reactive CD4^+^ T-cell responses to SARS-CoV-2 that were attributable to previous HCoV infections (34–37). The impact of this phenomenon on immune responses to vaccination against SARS-CoV-2 has to date only been investigated in few studies (33). Previous reports collectively suggest that pre-existing cross-reactive cellular immunity may boost mRNA vaccine-induced spike-specific CD4^+^ T-cell responses and neutralizing antibody levels against SARS-CoV-2 (33, 38–40). However, precise impacts on the kinetics of these responses are yet unclear.

It is well established in literature that memory T-cells generally produce more robust and rapid responses to secondary infection compared to naïve T-cells (41). Additionally, memory CD4^+^ T-cells provide accelerated help and promote significantly earlier class switching in primary B-cells compared to naïve CD4^+^ T-cells over the course of an infection (19). Accordingly, we hypothesize that individuals with pre-existing cross-reactive T-cell immunity to SARS-CoV-2 will exhibit boosts in vaccine-induced spike-specific T-cell responses and neutralizing antibody levels.

Investigating the precise relationship between the kinetics of vaccine-induced cellular and humoral responses to SARS-CoV-2, and how pre-existing cross-reactive immunity may impact these responses, is of paramount importance for informing the development of effective pan-coronavirus vaccines and vaccine regimens that can maximize the durability and breadth of protection against both current and future human coronaviruses of concern. To pursue our objectives, dual-colour ELISpot, ELISA, and live SARS-CoV-2 neutralization assays were deployed to longitudinally assess SARS-CoV-2 spike-specific IFN-γ and IL-2 T-cell responses, serological anti-spike/RBD IgG and IgA antibody levels, and neutralizing capacity in high-risk long-term care home (LTCH) staff up to 6-months post-second dose of BNT162b2 or mRNA-1273. Non-linear mixed-effects mathematical modelling was additionally deployed to precisely evaluate the kinetics of cellular and humoral responses to mRNA vaccination overtime.

## MATERIALS AND METHODS

### Study Cohort

From February to December 2021, peripheral blood mononuclear cell (PBMC) and serum samples were collected from 139 LTCH staff recruited from 12 LTCHs across the Greater Toronto Area and St. Champlain regions of Ontario, Canada. Samples were collected up to 6-months post-second dose of BNT162b2 or mRNA-1253. The LTCH staff recruited into the study were particularly those who worked in high-risk long-term care homes and resided in higher-density households within high-risk neighborhoods. By May 20^th^, 2020, LTCH staff in these regions comprised 27% of all active COVID-19 cases in Ontario (42, 43). The active case rate later dropped to 1.4% by January 12^th^, 2021, due to rapid vaccination efforts (42, 43). All sample acquisition was completed in compliance with a research ethics board (REB)-approved protocol (University of Toronto, Toronto, Canada; REB20-347).

### COVID-19 Screening of LTCH Staff

In accordance with the directive issued by the Minister of Long-Term Care of Ontario regarding LTCH SARS-CoV-2 surveillance testing and access to LTCHs, the LTCH staff cohort studied in this paper was required to take either:

a. One PCR test and one rapid antigen test (RAT) on separate days within a period of seven days. The period between PCR testing and RAT was required to be as close to seven days as practically achievable;
b. A RAT at least twice a week on separate days if fully vaccinated (at least 2 doses) against COVID-19; or
c. A RAT at least three times per week on separate days if not fully vaccinated against COVID-19

COVID-19 screening of LTCH staff was continuously conducted across all timepoints in this study.

### PBMC Isolation

PBMCs were isolated from whole blood samples collected in 8.5mL BD Vacutainer glass collection tubes with acid citrate dextrose (ACD) used as an anticoagulant (Fisher Scientific). Two tubes were collected from each participant. Whole blood was layered on Ficoll-Paque Plus (GE Healthcare) in 50mL SepMate tubes (Stemcell Technologies) and centrifuged at 1,200xg for 10 minutes. The buffy coat was then isolated and washed twice with 2% heat-inactivated fetal bovine serum (FBS; Wisent) in Dulbecco’s phosphate-buffered saline (D-PBS; Wisent) before being resuspended in R-10 medium. R-10 medium consisted of a mixture of RPMI 1640 (Wisent), 10% FBS (Wisent), 10mM HEPES (Wisent), 2mM L-glutamine (Wisent), and 100 UI/mL of penicillin-streptomycin (Wisent). Resuspended PBMCs were then diluted 1:1 with freezing medium consisting of 20% dimethyl sulfoxide (DMSO; Sigma-Aldrich) and 80% FBS and subsequently aliquoted for storage at −150 °C.

### Serum Isolation

Serum was isolated from whole blood collected in 3.5mL BD Vacutainer serum separation tubes containing spray-coated silica and a polymer gel for serum separation (Fisher Scientific). One tube was collected from each participant. Tubes were centrifuged at 1,200xg for 10 minutes before serum was subsequently isolated and aliquoted for storage at −80 °C.

### SARS-CoV-2 Peptide Masterpool Synthesis

SARS-CoV-2 15-mer peptides from nucleocapsid phosphoprotein (N), envelope glycoprotein (E), membrane glycoprotein (M), and non-structural protein (NSP) were synthesized by GeneScript (Piscataway, NJ) using the wild-type SARS-CoV-2 reference sequence (accession number NC_045512.2, NCBI). The SARS-CoV-2 spike glycoprotein masterpool was obtained as a PepMix (Swiss-Prot ID: P0DTC2) from JPT (Berlin, Germany). The N, E, M, and S masterpools were respectively comprised of 102, 12, 49, and 315 (split into two pools of 158 and 157) 15-mer peptides with 11 amino acid overlaps. Each of the four aforementioned masterpools spanned the entirety of their corresponding antigens. The NSP masterpool consisted of 26 15-mer peptides that were collectively derived from 14 non-structural antigens listed in **Table S1**. These peptides were previously reported to elicit cross-reactive responses toward SARS-CoV-2 in healthy, unexposed individuals (34). All peptides were reconstituted in 100% DMSO and aliquoted for storage at −80 °C until needed.

### *Ex Vivo* ELISpot Assay

To perform *Ex vivo* ELISpot assays, MSIPS4W plates (Milipore) were first activated with 35% ethanol. Activated plates were then washed with molecular grade sterile water (Wisent) before coating with 10 ug/mL of monoclonal IFN-γ (1-D1K) and 10ug/mL of monoclonal IL-2 (MT2A91/2C95) capture antibodies (Mabtech). The coated plates were wrapped in parafilm wax and allowed to incubate overnight at 4°C. Following incubation, coated plates were washed with D-PBS and blocked with R-10 medium for 1 hour at 37°C, 5% CO_2_. During this time, PBMCs were thawed and rested for at least 2 hours at 37°C, 5% CO_2_. Following incubation, plates were washed with D-PBS and PBMCs were plated at 250,000 cells per well. Over a period of 24 hours at 37°C, 5% CO_2_, PBMCs were then stimulated with N, E, M, S, or NSP masterpools at a concentration of 1 ug/mL. As positive controls to assess T-cell functionality, PBMCs were stimulated with CEF (NIH HIV Reagent Program, #ARP-9808) and CEFTA (Mabtech, #3617-1) peptide pools at 1ug/mL. As a positive control to assess the assay’s functionality, PBMCs were stimulated with *Staphylococcal* enterotoxin B (SEB; Sigma-Aldrich, #S4881) at 0.1 ug/mL. At the end of the incubation period, PBMCs were discarded, and plates were washed with 0.05% Tween 20 (BioShop) in D-PBS. Plates were subsequently coated with a cocktail of alkaline phosphatase (ALP)-conjugated monoclonal IFN-γ detection antibody (7-B6-1; Mabtech) at 1:500 dilution and biotinylated monoclonal IL-2 detection antibody (MT8G10; Mabtech) at 0.25 ug/mL in D-PBS. Plates were allowed to incubate for 2 hours at room temperature (RT) in the dark. After incubating, plates were washed again with 0.05% Tween 20 in D-PBS before streptavidin-horseradish peroxidase (HRP) conjugate was plated at 1:1000 dilution in D-PBS. After incubating for another hour at RT in the dark, plates were washed with molecular grade sterile water. Spots were lastly developed by first treating wells with Vector Blue developing reagent for ALP (Vector Laboratories) for 15 minutes at RT in the dark. Plates were subsequently washed with molecular grade sterile water before wells were treated with Vector NovaRed developing reagent for HRP (Vector Laboratories) for 8 minutes at RT in the dark. Plates were then washed a final time with MiliQ water and allowed to dry at 4◌° C in the dark. Spots in each well were quantified using an ImmunoSpot S3 Analyzer (Cellular Technology Limited).

To analyze the data, the average number of spots + 2 SD per million PBMC in negative control (DMSO) wells, or a minimum of 10 spots per million PBMC – whichever is greater – was subtracted from gross spot counts per million PBMC in treatment wells. Resulting net spot counts greater than 0 were considered positive.

### *In Vitro* PBMC Expansion

To expand PBMCs *in vitro*, cryopreserved PBMCs were thawed and washed twice with R-H10 medium consisting of a mixture of RPMI 1640 (Wisent), 10% heat-inactivated human AB serum (Wisent), 10mM HEPES (Wisent), 2mM GlutaMAX^TM^ (Thermo Fisher Scientific), 100 UI/mL of penicillin-streptomycin (Wisent), 1mM sodium pyruvate (Thermo Fisher Scientific), and 50uM 2-mercaptoethanol (Thermo Fisher Scientific). PBMCs were adjusted to a final concentration of 2×10^6^ PBMC/mL before being plated at 400,000 cells (200uL) per well per condition within a tissue-culture treated 96-well round-bottom polystyrene plate. After plating, PBMCs were treated with 0.1% DMSO (negative control) or with 0.1μg/mL of NSP, CEF, CEFTA, or SEB and incubated at 37°C, 5% CO_2_ for a total of 7 days. On day 1 of incubation (with cell plating being day 0), PBMCs treated with 0.1% DMSO or 0.1ug/mL of NSP, CEF, or CEFTA were supplemented with 10IU/mL of recombinant human IL-2 (rh IL-2; R&D Systems) and 10ng/mL of recombinant human IL-7 (rh IL-7; R&D Systems). PBMCs treated with 0.1ug/mL of SEB were supplemented with 10ng/mL of rh IL-7 alone. By day 5 of incubation, half the culture medium in each well was carefully aspirated and replaced with fresh R-H10 medium. DMSO, NSP, CEF, and CEFTA wells were additionally re-supplemented with 10IU/mL of rh IL-2 and 10ng/mL of rh IL-7, while SEB wells were re-supplemented with 10ng/mL of rh IL-7 alone. Lastly, on day 7, PBMCs were washed twice with R-10 medium after centrifuging at 300xg for 5 minutes before being assayed by ELISpot.

### Chemiluminescent ELISA Assay for Detection of SARS-CoV-2-Specific IgG and IgA Antibodies in Serum

A chemiluminescent ELISA assay was used to measure levels of serological IgG and IgA antibody to the full-length spike trimer, the receptor binding domain (RBD), and nucleocapsid of the ancestral SARS-CoV-2 virus, as previously described (44, 45). Briefly, LUMITRAC 600 high-binding white polystyrene 384-well microplates (Greiner Bio-One, #781074; VWR, #82051-268) were first pre-coated overnight with 10uL per well of antigen (Ag): 50 ng of full-length spike trimer (SmT1), 20ng of RBD (331–521), and 7ng of nucleocapsid. All antigens were supplied by the National Research Council of Canada (NRC). Following overnight incubation, microplates were washed 4 times at RT with 100uL per well of PBS-T before each of the following steps. Step one: wells were blocked for 1 hour in 80uL of 5% Blocker BLOTTO (ThermoFisher Scientific, #37530). Step two: 10uL of serum diluted to 1:160, 1:640, 1:2560, or 1:40960 with 1% Blocker BLOTTO in PBS-T was added to each well and incubated for 2 hours. Step three: 10uL of HRP-fused human anti-IgG (IgG#5 by NRC, 0.9 ng/well) or HRP-conjugated human anti-IgA (Jackson ImmunoResearch, #109-035-127, 0.8 ng/well) diluted with 1% Blocker BLOTTO in PBS-T was added to each well, followed by a 1-hour incubation. Step four: 10uL of ELISA Pico Chemiluminescent Substrate (ThermoFisher Scientific, #37069, diluted 1:4 in MilliQ distilled H_2_O) was added and incubated for 5-8 minutes. Chemiluminescence was read on an EnVision 2105 Multimode Plate Reader (Perkin Elmer) at 100 ms/well using an ultra-sensitive detector. Raw chemiluminescent values were normalized to a synthetic standard included on each assay plate (for IgG: VHH72-Fc supplied by NRC for spike/RBD or an anti-nucleocapsid IgG Ab from Genescript, #A02039; for IgA: anti-spike CR3022 from Absolute Antibody, #Ab01680-16.0 and anti-nucleocapsid CR3018 from Absolute Antibody, #Ab01690-16.0). Computed relative ratios were further converted to binding Ab units (BAU/mL) using the WHO International Standard 20/136 as the calibrant (44). A positivity threshold for the 1:160 dilution was established as being 3SD from the mean of negative control samples, as previously described (44).

All measures of serological antibody levels that did not meet positivity thresholds were plotted as 0.

### Chemiluminescent Direct ELISAs for HCoV Spike-Specific IgG Antibody Detection in Serum

Automated chemiluminescent ELISAs were based upon and optimized from assays first described in (46). Assays were performed using Hamilton Microlab STAR robotic liquid handlers at the University of Ottawa’s Serology and Diagnostics High-Throughput Facility (Faculty of Medicine). To prepare assay plates, antigens (Spike trimers from seasonal coronavirus strains: 229E (DAGC134), HKU1 (DAGC132), OC43 (DAGC131) and NL63(DAGC133); Creative Diagnostics), were diluted in PBS and applied to wells of a 384-well high-binding polystyrene Nunc plate (Thermo Fisher Scientific, #460372) at a final amount of 50 ng/well. Antigen-coated plates were centrifuged briefly at approximately 2,000xg in a plate spinner (Fisher Scientific) to ensure even coating then incubated on a rocker at 4°C (minimum 4 hours, up to 3 days). On the day of the assay, antigen-coated plates were washed, and wells were blocked with 80 µL of 3% w/v skim milk powder dissolved in PBS + 0.1% Tween (PBST) for 1 h. Samples and controls were diluted at 1:100, 1:1000 and 1:10000 in 1% w/v skim milk powder in PBST in a 96-well deep-well plate. Following the blocking incubation, plates were washed, and 10 µL of diluted samples and controls were added to their respective wells. Plates were incubated for 2 h and wells were then washed. Anti-IgG (NRC anti-hIgG#5-HRP fusion) was diluted at 1:5400 in 1% w/v skim milk powder in PBST and 10 µL was added to each well and incubated for 1 h. Plates were washed and 10 µL of ELISA Pico Chemiluminescent Substrate (Thermo) diluted 1:2 in MilliQ H_2_O, was dispensed into each well. After a subsequent 8-minute incubation period, plates were read on a Synergy Neo2 plate reader (BioTek Instruments) at 20 ms/well and a read height of 1.0 mm. All incubations apart from antigen coating were performed at room temperature, and all plate washes were conducted using a 405 TS/LS LHC2 plate washer (Biotek Instruments); all wash steps included four washes with 100 μL PBST.

Blank-subtracted raw luminescence reads were normalized between dilutions by applying a scaling factor derived from the average ratios of the medians of on-plate control medians. On-plate controls at 1:1000 dilution were used as the reference point for normalization.

### Live Ancestral SARS-CoV-2 Neutralization Assay

Neutralizing capacities of LTCH staff sera were assessed using live ancestral SARS-CoV-2 neutralization assays. LTCH staff serum samples were heat inactivated in a hot water bath at 56°C for 30 minutes and then allowed to cool to RT prior to use in live ancestral SARS-CoV-2 neutralization assays. All assays were conducted using VeroE6 cells (ATCC #CRL-1586) cultured in D-10 medium (DMEM supplemented with 10% heat inactivated FBS, 100U/mL Penicillin, 100U/mL Streptomycin, and 2mM L-Glutamine). Briefly, 6 x 10^4^ VeroE6 cells were seeded per well of a 96-well flat bottom tissue culture plate and rested overnight at 37°C, 5% CO_2_. LTCH staff serum samples were then serially diluted in serum-free DMEM and incubated in the presence of 100TCID_50_ of ancestral SARS-CoV-2 (SARS-CoV-2-SB2-PB clone 1) for 1 hour at 37°C, 5% CO_2_. Following incubation, VeroE6 cells were inoculated with the virus/serum co-culture for 1 hour at 37°C, 5% CO_2_. The inoculum was subsequently removed from all wells and replaced with D-2 medium (DMEM supplemented with 2% heat inactivated FBS, 100U/mL Penicillin, 100U/mL Streptomycin, and 2mM L-Glutamine) before being incubated for 5 days at 37°C, 5% CO_2_. Following the incubation period, the cytopathic effect (CPE) of the viral inoculation was visually assessed by examining VeroE6 cells under a phase-contrast microscope. All live ancestral SARS-CoV-2 neutralization assays were conducted in quadruplicates. Results were analyzed using PRISM GraphPad software (GraphPad Software, La Jolla, Ca). A 4-paramater logistic non-linear regression analysis was performed to compute IC_50_ values, which represent serum concentrations at which 50% of wells were negative for CPE. All handling of the live ancestral SARS-CoV-2 virus was performed within the Combined Containment Level 3 (C-CL3) Unit at the Temerty Faculty of Medicine, University of Toronto.

All values of serological neutralizing capacity are provided as log(1/IC50). Sera with no neutralizing capacity were assigned a value of 0.

### Mathematical Modelling Approach and Parameter Estimation

A mathematical model of exponential growth (eq. 1) or decay (eq. 2) was fit to individual study data. All fits were performed in Monolix (Version 2020R1) using non-linear mixed-effects models. For each data type (e.g. IFN-γ), we simultaneously fit all individuals to determine best-fit individual responses as well as a best-fit population response. Exponential growth/decay rates were determined for patients with increasing/decreasing measures of IFN-γ, IL-2, IgG, IgA, and log(1/IC50). For each fit we computed random effect parameters (and relative standard error on random effects). Random effects are practical identifiability analyses to characterize the data-driven model identifiability outcomes. We further compute and provide the relative standard error on each set of fitted population parameters. The equations for growth and decay kinetics are,

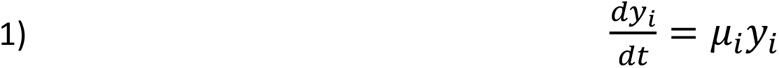

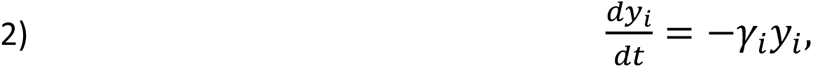

where *i* corresponds to the *i ^th^* individual in the study, *y* corresponds to the clinical feature being fit (i.e. IFN-γ, IL-2, IgG, IgA, or log[1/IC50]), *μ*_i_is the growth rate and *γi* is the decay rate.

### Statistical Analysis

The non-parametric Kruskal-Wallis Test (47) was employed to compute cross-sectional comparisons at similar timepoints between groups within categorizations of sex and vaccine types. The non-parametric Dunn’s test was employed to compute cross-sectional comparisons at similar timepoints between n = 5 immunological phenotype groups. P-values were corrected for multiples comparisons using the Benjamini-Hochberg method. The non-parametric Wilcoxin signed-rank test was employed to compute longitudinal comparisons from 2-6 weeks to 6-months post-second dose of BTN162b2 or mRNA-1273. * = 0.01 < p ≤ 0.05, ** = 0.001 < p ≤ 0.01, *** = 0.0001 < p ≤ 0.001, **** = p ≤ 0.0001. All correlations between spike-specific T-cell responses to mRNA vaccination and serological anti-spike/RBD IgG and IgA antibody levels or neutralizing capacity were computed using non-parametric Spearman tests (r_s_ = .00-.19 [very weak]; r_s_ = .20-.39 [weak]; r_s_ = .40-.59 [moderate]; r_s_ = .60-.79 [strong]; r_s_ = .80-1.0 [very strong]).

## RESULTS

### Sample Acquisition

To investigate how SARS-CoV-2-specific T-cell responses contribute to mRNA vaccine-induced humoral immunity, T-cell and humoral responses to BNT162b2 or mRNA-1273 were longitudinally assessed in a cohort of 139 high-risk LTCH staff. PBMC and serum samples were collected across three timepoints from February 10^th^ to December 14^th^, 2021: baseline (up to 7-days post-first dose, n = 10; low sample size due to rapid vaccine rollouts and delays in obtaining REB approval); 2-6 weeks post-second dose of BNT162b2 or mRNA-1273 (n = 137, 129 were additionally enrolled, 2 withdrew from baseline); and 6-months post-second dose (n = 117, 20 withdrew or were not analyzed because they received a third dose of mRNA vaccine). Refer to **Figure 1** for a graphical overview of the LTCH staff cohort throughout the study. According to Ontario public health records, the Alpha variant of SARS-CoV-2 predominated from February to late May 2021. Delta later became the major variant of concern from June to December 2021 (48–50).

**Figure 1:**
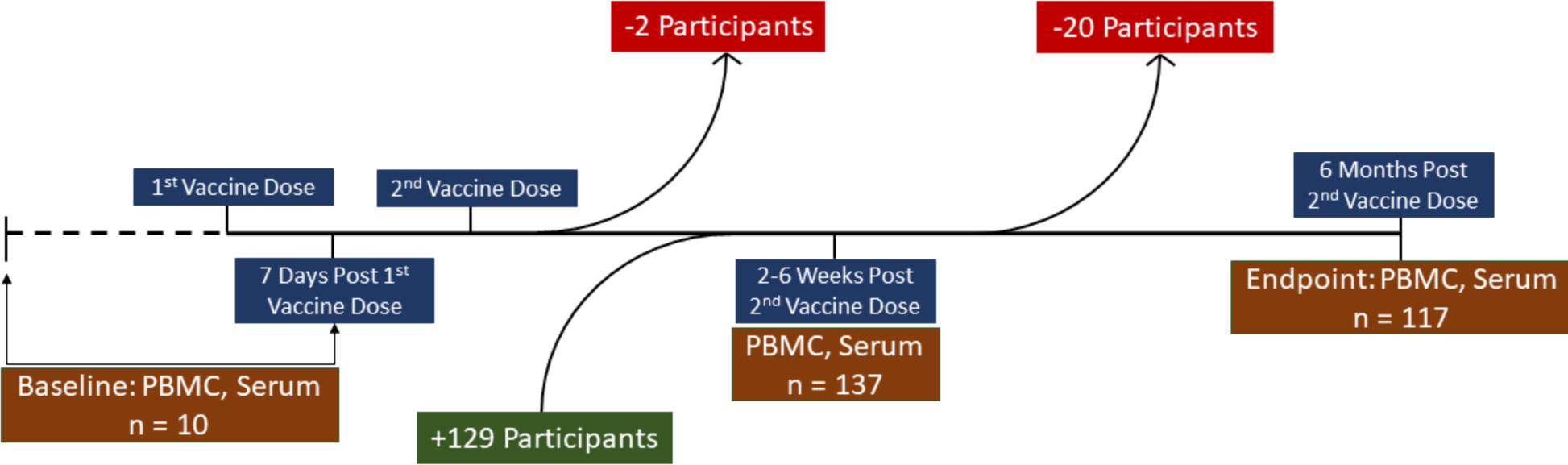
An overview of the LTCH staff cohort throughout the study.

### Immunological Phenotypes of LTCH Staff

To assess the effects of SARS-CoV-2-specific T-cell immunity on mRNA vaccine-induced humoral immunity, the following clinical features were used to stratify participants into separate immunological phenotypes:

1. History of previous PCR- or RAT-confirmed SARS-CoV-2 infections (see methods);
2. T-cell responses against SARS-CoV-2 N, E, M, S, and NSP masterpools by ELISpot at baseline or 2-6 weeks post-second dose; and
3. Anti-nucleocapsid (anti-N) IgG/IgA antibody levels in the serum.

Based on these clinical features, a total of 5 immunological phenotypes were defined as follows:

1. **Uninfected participants:** Participants who consistently tested negative for SARS-CoV-2 by PCR/RAT and remained seronegative for anti-N IgG/IgA antibodies both prior to and during the study were denoted as ‘uninfected’. Uninfected participants could be further sub-stratified into two subgroups:

a. **Non-T-cell Cross-Reactive (NCR):** Those who failed to exhibit detectable IFN-γ or IL-2 T-cell responses by ELISpot to any SARS-CoV-2 masterpools at baseline or to any non-spike masterpools at 2-6 weeks post-second dose (**Figure 2**).
b. **T-cell Cross-reactive (CR):** Those who demonstrated detectable IFN-γ or IL-2 T-cell responses by ELISpot to any SARS-CoV-2 masterpool at baseline or to any non-spike masterpools at 2-6 weeks post-second dose (**Figure 2**).
2. **Hybrid Immune (HI):** Participants who contracted a PCR/RAT-confirmed SARS-CoV-2 infection prior to either the first or second dose of BNT1262b2 or mRNA-1273. Refer to **Table S2** for a summary of clinical characteristics.
3. **Asymptomatic Breakthrough:** Participants who seroconverted to possess anti-N IgG/IgA antibodies between 2-6 weeks and 6-months post-second dose without ever displaying COVID-19-related symptoms or testing positive by PCR/RAT. Refer to **Table S3** for a summary of clinical characteristics. No cases of symptomatic breakthrough infections were detected.
4. **Anti-N+, PCR**(**-**): Participants who were already seropositive for anti-N IgG/IgA antibodies upon initial sample collection with no prior history of COVID-19-related symptoms or PCR/RAT-confirmed SARS-CoV-2 infections. It has been shown by previous literature that 20-23% of pre-pandemic individuals – from as early as 2014 – possessed serological titers of anti-N antibodies that were cross-reactive with SARS-CoV-2 (51, 52). Accordingly, since it was not possible to determine whether N+, PCR(-) participants identified at initial sample collection were convalescent or healthy with pre-existing humoral cross-reactivity, they were decidedly stratified into their own group.
5. **Unknown:** Participants who could not be clearly categorized due to an absence of cellular or serological data. These individuals could not be included in any analyses.

**Figure 2:**
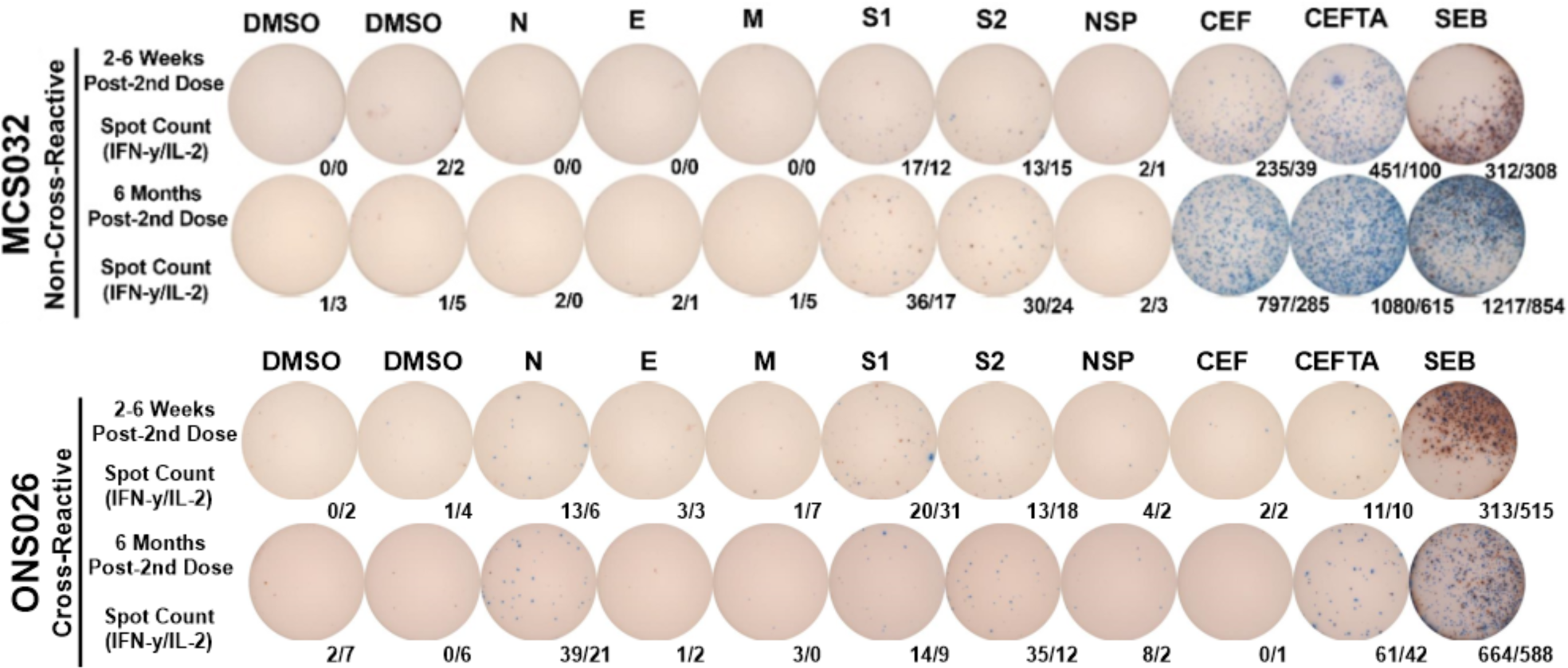
A representative ELISpot comparing spike and non-spike-specific IFN-γ (blue) and IL-2 (red) T-cell responses between non-cross-reactive and cross-reactive vaccinees. Spot counts are provided as IFN-γ/IL-2 spot-forming cells (SFC) per 250,000 PBMC. DMSO = negative controls; N = nucleocapsid; E = envelope; M = membrane; S1+S2 = full spike; NSP = non-structural protein; CEF+CEFTA+SEB = positive controls.

**Table 1** provides a summary of the clinical characteristics of the LTCH staff cohort at each timepoint of the study.

**Table 1:**
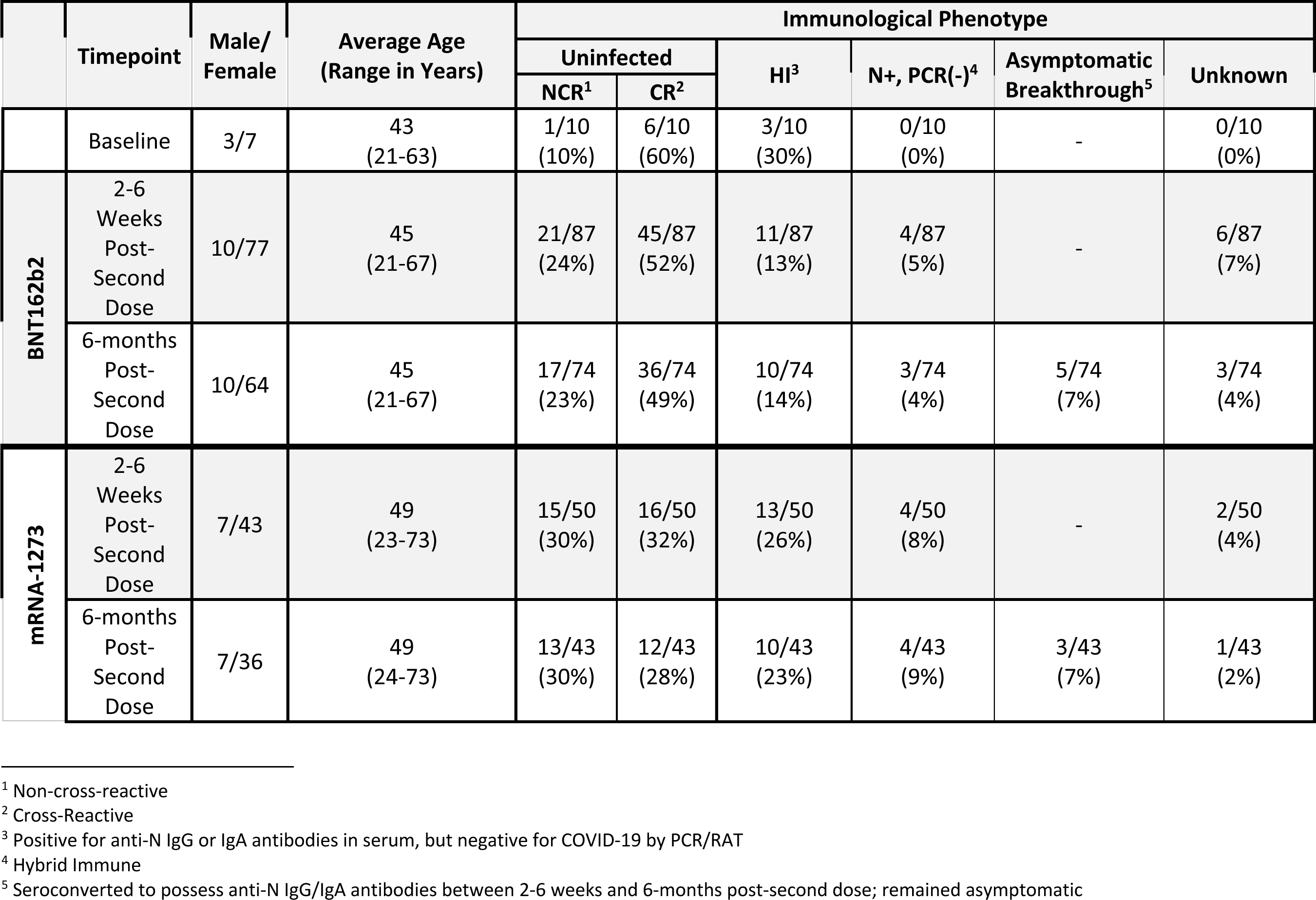
A summary of clinical characteristics of the LTCH staff cohort at each timepoint.

### Comparing HCoV Antibody Levels between CR and NCR Vaccine Recipients

Given former evidence of the induction of cross-reactive T-cell immunity to SARS-CoV-2 by previous HCoV infections (34–37), we sought to determine whether CR vaccinees possessed elevated serological anti-spike IgG antibody levels against HCoV-HKU1, HCoV-NL63, HCoV-OC43, or HCoV-229E compared to NCR vaccine recipients at 2-6 weeks post-second dose. Antibody levels were observed to be highly similar between the two groups (**Figure 3**).

**Figure 3:**
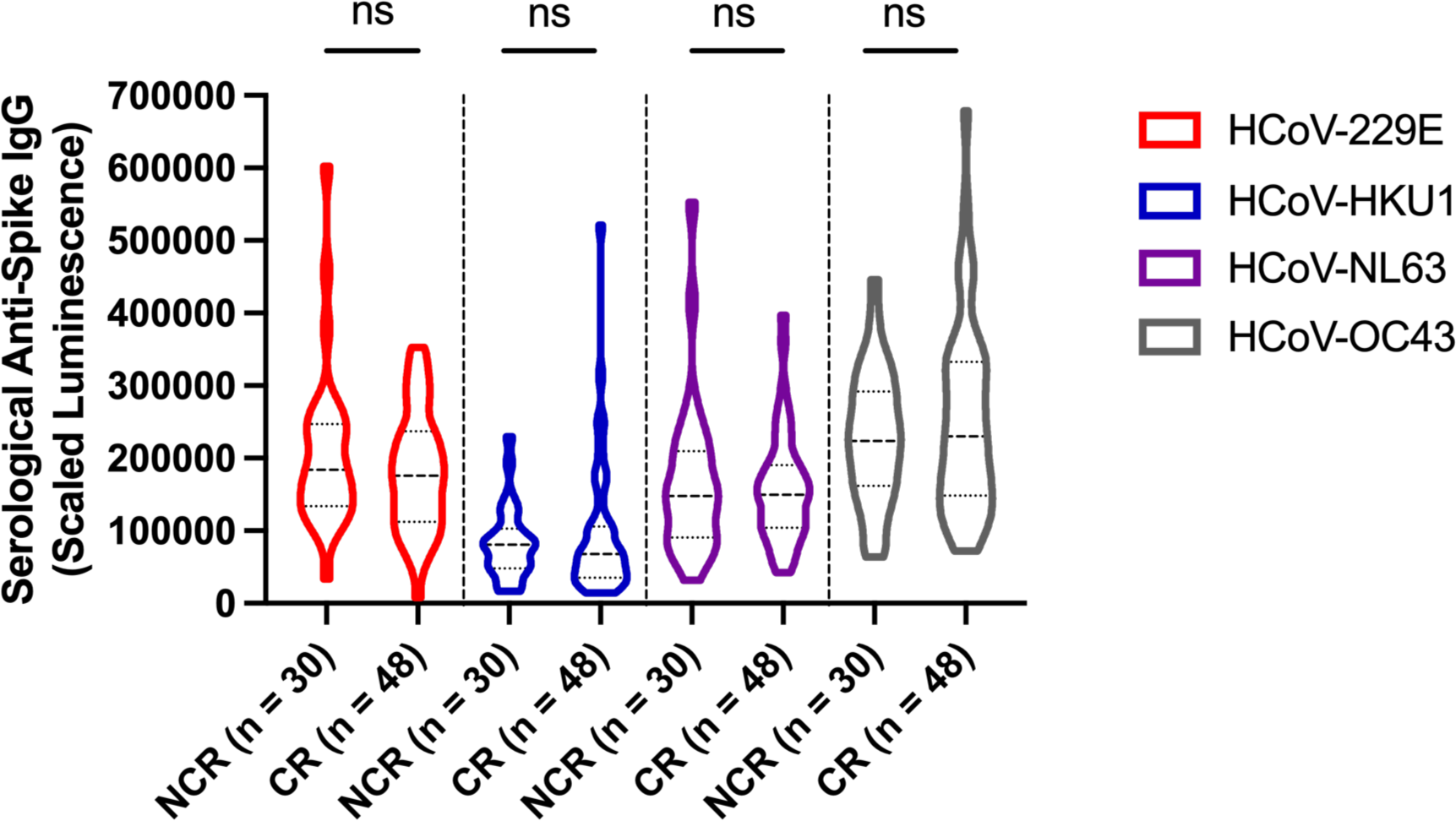
Comparisons of serological anti-spike IgG antibody levels against HCoV-229E (red), HCoV-HKU1 (blue), HCoV-NL63 (purple), and HCoV-OC43 (grey) between non-cross-reactive and cross-reactive mRNA vaccinees at 2-6 weeks post-second dose.

### Comparing SARS-CoV-2 Non-Spike-Specific T-cell Responses between CR and HI Vaccine Recipients

As reported in previous studies, pre-existing cross-reactive T-cells in unexposed individuals frequently target regions of SARS-CoV-2 structural and non-structural antigens that are conserved between SARS-CoV-2 and the endemic HCoVs (34, 35, 37, 40, 53, 54). We thus investigated how the distribution of T-cell responses across non-spike antigens of SARS-CoV-2 compared between CR and HI vaccine recipients following the second dose of BNT162b2 or mRNA-1273. Anti-N+, PCR(-) (n = 7) participants were not be included in these analyses due to a very small sample size.

As shown in **Figure 4A**, HI vaccine recipients mounted significantly greater median dual IFN-γ/IL-2 T-cell responses against M compared to CR recipients at 2-6 weeks post-second dose (p = 0.007). Although differences were later lost by 6-months post-second dose (**Figure 4A**), M-specific T-cell responses began to dominate the total non-spike-specific response in HI vaccine recipients (**Figure 4B**). CR recipients contrarily began to mount the majority of non-spike specific T-cell responses against the NSP masterpool (**Figure 4B**), which is comprised of 15mer peptides with a considerable degree of homology with endemic HCoVs (34).

**Figure 4:**
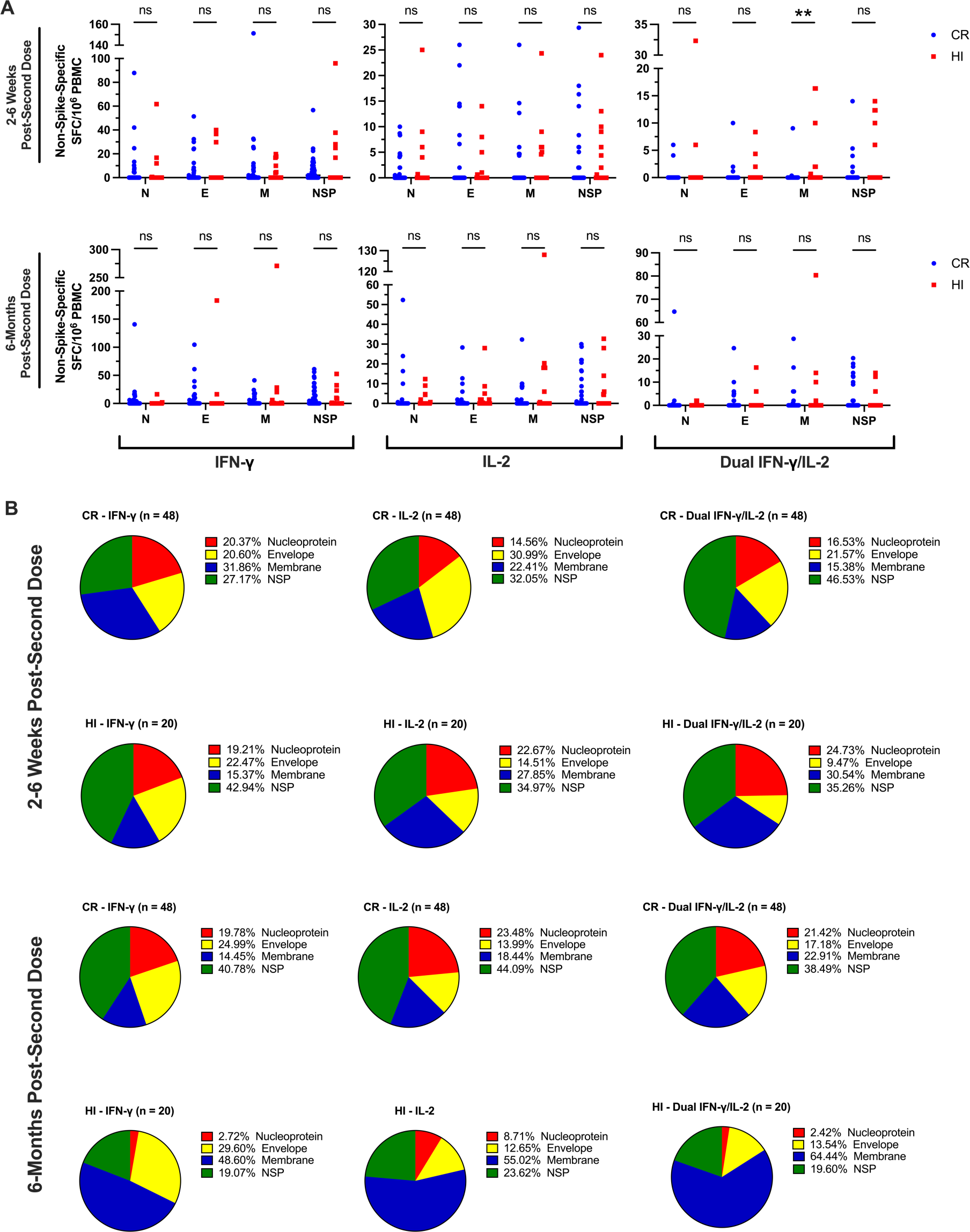
(A) Scatterplot comparisons of N-, E-, M-, and NSP-specific T-cell responses between CR (blue circles) and HI (red squares) vaccine recipients. (B) Pie chart comparisons of the distribution of IFN-γ, IL-2, and dual IFN-γ/IL-2 T-cell responses to SARS-CoV-2 nucleoprotein (red), envelope (yellow), membrane (blue), and NSP (green) peptide masterpools between HI and CR vaccine recipients of BNT162b2 or mRNA-1273 at (A) 2-6 weeks and (B) 6-months post-second dose.

### Assessment of Ancestral SARS-CoV-2 Spike-Specific T-cell Responses in LTCH Staff over the Course of 6-months Post-Second Dose of BNT162b2 or mRNA-1273

We next studied mRNA vaccine-induce spike-specific T-cell responses in LTCH staff following mRNA vaccination with BNT162b2 or mRNA-1273. Dual-colour IFN-γ/IL-2 ELISpot assays were used to characterize these responses at 2-6 weeks and 6-months post-second dose. Mathematical models of exponential growth/decay were additionally fit to individual datapoints to analyze the precise kinetics of spike-specific T-cell responses across the two timepoints. Due to a scarcity of sample, ELISpot assays could only be conducted on whole PBMCs without any enrichment for the CD4^+^ T-cell subset.

Relative to baseline, robust spike-specific IFN-γ, IL-2, and dual IFN-γ/IL-2 T-cell responses were observed in LTCH staff at 2-6 weeks post-second dose of BNT162b2 or mRNA-1273 (**Figure 5A-C; Table S4**). Over the course of 6-months post-second dose, 61/113 (54%) LTCH staff analyzed unexpectedly exhibited a growth in spike-specific IFN-γ T-cell responses with an average doubling time of 173 (150–203) days (**Figure 6A-C**). The remaining 52/113 (46%) had experienced a decay with an average half-life of 115 (69–346) days (**Figure 6A-C**). With respect to spike-specific IL-2 T-cell responses, 48/113 (42%) LTCH staff likewise exhibited a growth with an average doubling time of 173 (99–693) days (**Figure 6A, B, D**). The remaining 65/113 (58%) experienced a decay with an average half-life of 115 (63–693) days (**Figure 6A, B, D**). Resultantly, spike-specific IFN-γ and IL-2 T-cell responses across the entire cohort at 6-months post-second dose were not observed to significantly differ from those observed at 2-6 weeks post-second dose (**Figure 5A-C**).

**Figure 5:**
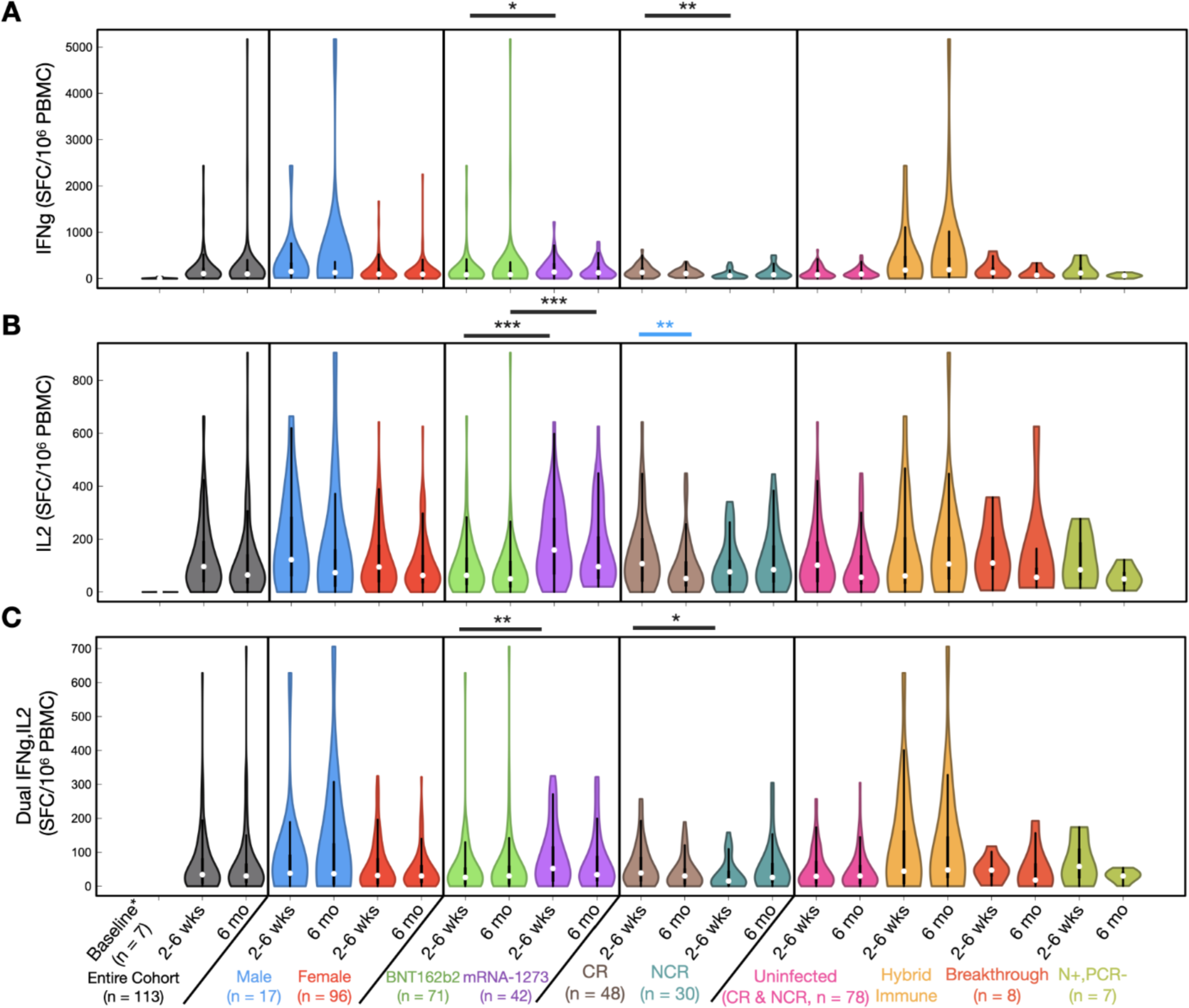
(A-C) comparisons of net spike-specific (A) IFN-γ, (B) IL-2, and (C) dual IFN-γ/IL-2 T-cell responses at 2-6 weeks and 6-months post-second dose. Squares and whiskers within each violin plot represent medians with interquartile ranges (IQRs). Refer to **Table S4** for medians with IQRs for each group. Significant cross-sectional comparisons are indicated with black asterisks. Significant longitudinal comparisons are indicated with blue asterisks. Baseline is comprised of a group of 7 uninfected LTCH staff.

**Figure 6:**
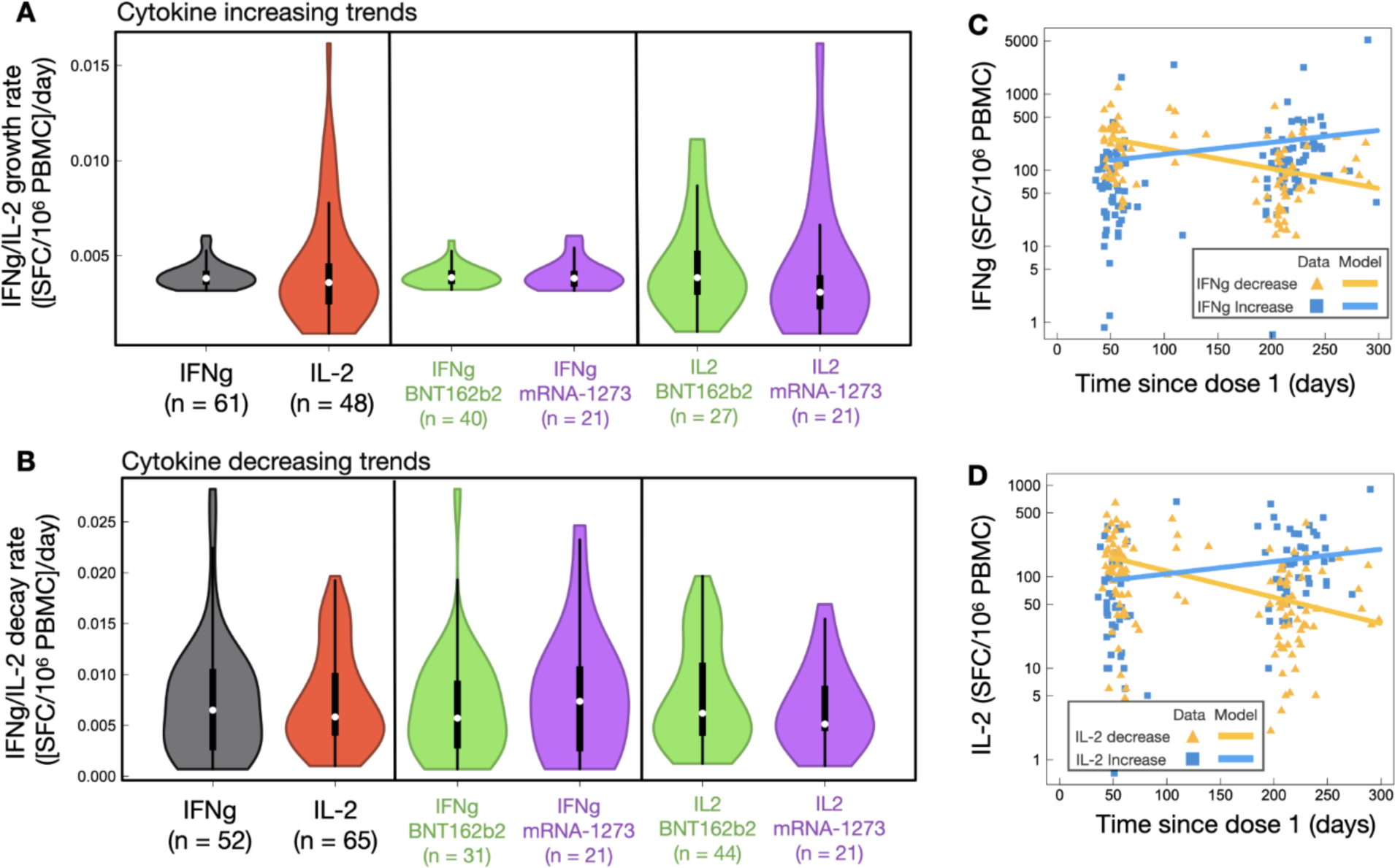
(A, B) Violin plot analysis of (A) growth and (B) decay rates in spike-specific IFN-γ and IL-2 T-cell responses across the entire cohort and between mRNA vaccine types. Squares and whiskers within each violin plot represent medians with IQRs. (C, D) Spike-specific (C) IFN-γ and (D) IL-2 T-cell responses as a function of time since dose 1 for n = 113 participants that could be analyzed longitudinally, separated by increasing (blue) and decreasing (yellow) trends. Refer to **Table S5** for a summary of median decay and growth rates with half-lives and doubling times.

Stratifying for vaccine type showed that recipients of mRNA-1273 generally exhibited significantly greater spike-specific T-cell responses than those of BNT162b2 at 2-6 weeks post-second dose (**Figure 5B, C**). However, the durability of the response – defined by the rate of decline overtime – was not impacted by the mRNA vaccine platform. Stratifying for sex additionally did not uncover any differences in vaccine-induced spike-specific T-cell responses (**Figure 5A-C**) or durability (data not shown) between males and females.

To assess the influence of pre-existing T-cell cross-reactivity to SARS-CoV-2 on mRNA vaccine-induced spike-specific T-cell responses, uninfected participants were stratified into NCR and CR vaccine recipients. In comparison to NCR vaccinees, CR recipients of BNT162b2 or mRNA-1273 exhibited a significant boost in spike-specific IFN-γ (p = 0.0014) and dual IFN-γ/IL-2 (p = 0.034) T-cell responses at 2-6 weeks post-second dose (**Figure 5A, C**). However, these boosts were transient and observed to dissipate by 6-months post-second dose (**Figure 5A, C**). CR vaccine recipients additionally experienced significant declines (p = 0.0051) in spike-specific IL-2 T-cell responses by 6-months post-second dose, which were contrastingly entirely preserved in NCR vaccine recipients (**Figure 5B**).

No significant differences in spike-specific T-cell responses could be detected between uninfected participants and all other immunological phenotypes.

### Assessment of Serological Anti-Spike and Anti-RBD IgG and IgA Antibody Levels over a Period of 6-months Post-Second Dose of BNT162b2 or mRNA-1273

ELISA assays measuring levels of anti-spike and anti-RBD IgG and IgA antibody levels in sera were deployed to characterize humoral responses induced by mRNA vaccination in LTCH staff at 2-6 weeks and 6-months post-second dose. As before, mathematical models of exponential grow/decay were fit to individual data points for the analysis of response kinetics across the two timepoints.

Relative to baseline, robust anti-spike and anti-RBD IgG antibody levels were observed across the entire LTCH staff cohort at 2-6 weeks post-second dose of BNT162b2 or mRNA-1273 (**Figure 7A, B**; **Table S6**). However, over the course of 6-months post-second dose, 111/113 (98.5%) LTCH staff exhibited highly significant decays in anti-spike/RBD IgG antibody levels with an approximate half-life of 53 (50–58) days (**Figure 7A-F**).

**Figure 7:**
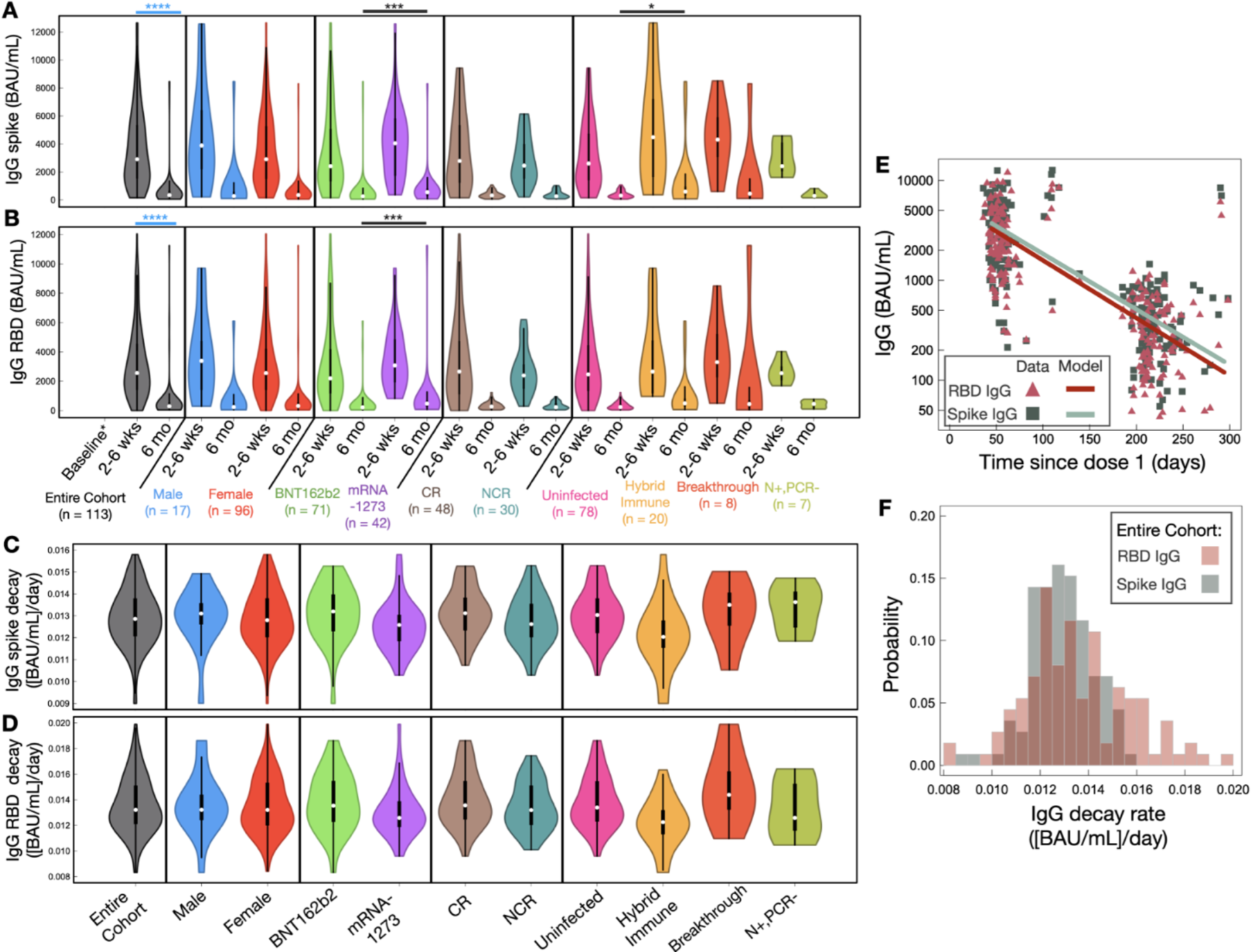
(A, B) violin plot analysis of serological (A) anti-spike IgG and (B) anti-RBD IgG antibody levels at 2-6 weeks and 6-months post-second dose. Squares and whiskers within each violin plot represent medians with IQRs (see **Table S6** for values). Significant cross-sectional comparisons are indicated with black asterisks. Significant longitudinal comparisons are displayed with blue asterisks for the entire cohort alone to avoid overcrowding the figure. Baseline is comprised of a group of 7 uninfected LTCH staff. (C, D) Violin plot analysis of (C) anti-spike IgG and (D) anti-RBD IgG decay rates across the entire LTCH staff cohort after adjusting for sex, vaccine type, and immunological phenotype. (E) Mathematical modelling of decay rates in anti-spike IgG (green) and anti-RBD IgG (red) antibody levels since dose 1 of BNT162b2 or mRNA-1273. (F) A histogram of individual decay rates in anti-spike IgG (green) and anti-RBD IgG (red) antibody levels. Refer to **Table S7** for a summary of median decay rates with half-lives for anti-spike and anti-RBD IgG antibody levels.

Compared to recipients of BNT162b2, anti-spike (p = 0.0613) and anti-RBD (p = 0.0704) IgG antibody levels were only slightly elevated in mRNA-1273 vaccinees at 2-6 weeks post-second dose (**Figure 7A, B**). However, in light of similar decay kinetics (**Figure 7C, D**), mRNA-1273 vaccinees later exhibited significantly greater anti-spike (p = 0.0003) and anti-RBD (p = 0.0009) IgG antibody levels at 6-months post-second dose (**Figure 7A, B**).

Stratifying for sex did not uncover differences in the magnitude or durability of anti-spike/RBD IgG antibody levels between males and females (**Figure 7A-D**).

Pre-existing cross-reactive T-cell immunity to SARS-CoV-2 was not observed to significantly boost mRNA vaccine-induced anti-spike or anti-RBD IgG antibody levels in uninfected vaccine recipients, although a trend was noted (**Figure 7A, B**). As shown in **Figures 7C, D** and **Table S7**, the durability of anti-spike and anti-RBD IgG antibodies were also highly similar between NCR and CR vaccine recipients and between all other immunological phenotypes.

With respect to anti-spike and anti-RBD IgA antibody levels, all LTCH staff possessed robust levels in the serum at 2-6 weeks post-second dose (**Figure 8A, B**; **Table S6**). As with IgG, 98.5% of participants exhibited a significant decline in anti-spike IgA antibody levels over the course of 6-months post-second dose with an approximate half-life of 76 (68–85) days (**Figure 8C-F**). Anti-RBD IgA antibody levels comparatively exhibited heightened median decay rates, translating to a reduced half-life of 59 (55–63) days (**Figure 8C-F**). Of importance to note, a 2-6-week window post-second dose may not have captured the initial rapid decay of IgA antibodies and may have thus overestimated half-life measures compared to IgG (26, 55–57).

**Figure 8:**
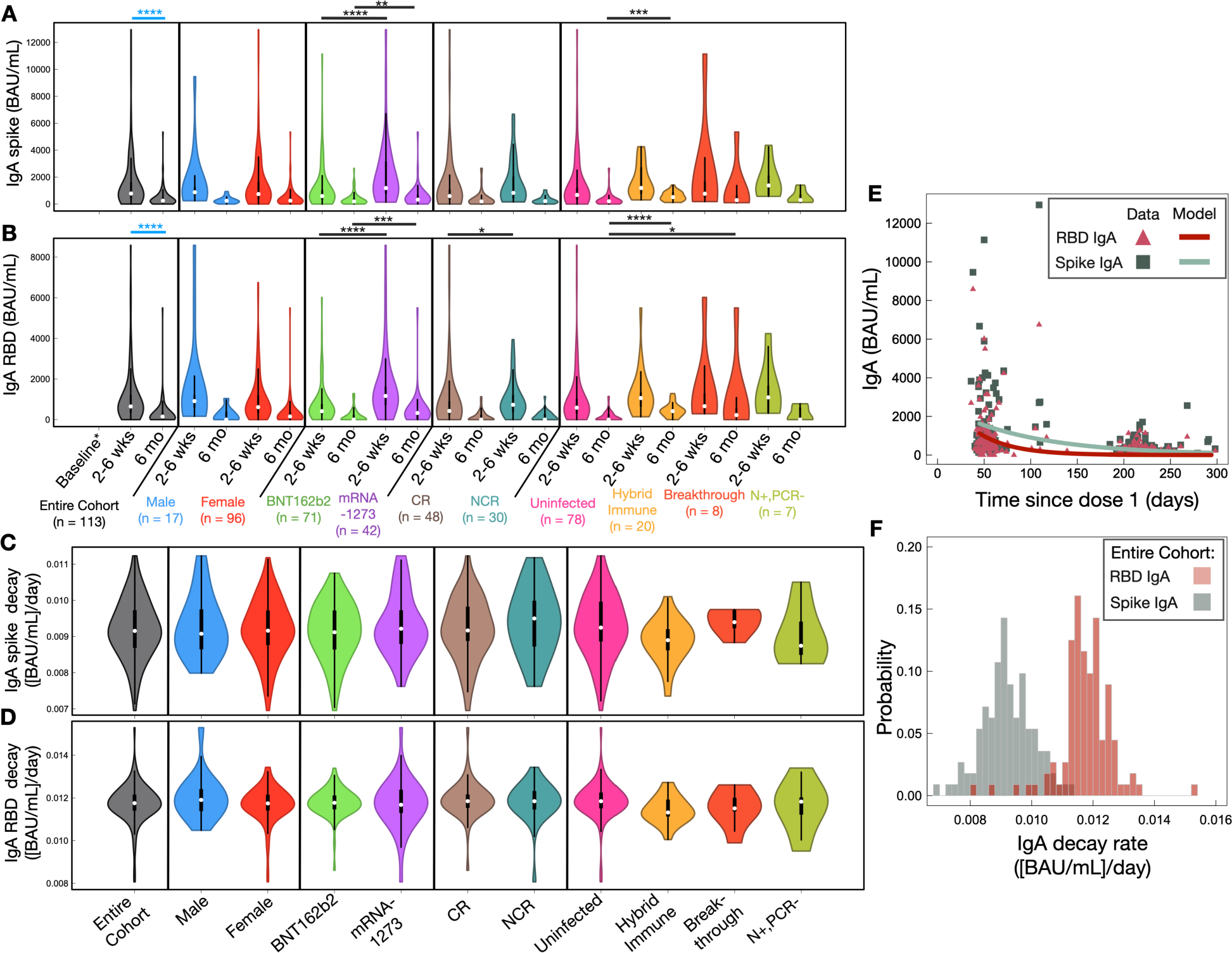
(A, B) violin plot analysis of (A) anti-spike IgA and (B) anti-RBD IgA antibody levels at 2-6 weeks and 6-months post-second dose. Squares and whiskers within each violin plot represent medians with IQRs. Refer to **Table S6** for medians with IQRs for each group. Significant cross-sectional comparisons are indicated with black asterisks. Significant longitudinal comparisons are displayed with blue asterisks for the entire cohort alone to avoid overcrowding the figure. Baseline is comprised of a group of 7 uninfected LTCH staff. (C, D) Violin plot analysis of serological (C) anti-spike IgA and (D) anti-RBD IgA decay rates across the entire LTCH staff cohort after adjusting for sex, vaccine type, and immunological phenotype. (E) Mathematical modelling of decay rates in anti-spike IgA (green) and anti-RBD IgA (red) antibody levels since dose 1 of BNT162b2 or mRNA-1273. (F) A histogram of individual decay rates in anti-spike IgA (green) and anti-RBD IgA (red) antibody levels (see **Table S7** for values).

Stratifying by vaccine type showed that recipients of mRNA-1273 had significantly higher anti-spike and anti-RBD IgA antibody levels than those of BNT162b2 recipients (p < 0.0001) at 2-6 weeks post-second dose (**Figure 8A, B**). Given that decay rates were highly similar between the two groups (**Figure 8C, D**; **Table S7**), anti-spike (p = 0.004) and anti-RBD (p = 0.0002) IgA antibody levels remained greater in mRNA-1273 vaccinees at 6-months post-second dose.

As before, sex was not observed to influence the magnitude or durability of spike- and RBD-specific IgA antibody levels (**Figure 8A-D**).

When stratifying our cohort by immunological phenotype, NCR vaccine recipients were observed to possess significantly greater anti-RBD IgA antibody levels (p = 0.044) than CR recipients at 2-6 weeks post-second dose (**Figure 8B**). However, differences between the two groups were transient and dissipated by 6-months post-second dose (p = 0.37; **Figure 8B**). No differences in anti-spike IgA antibody levels were observed between NCR and CR vaccine recipients at either timepoint (**Figure 8A**).

With respect to all other immunological phenotype groups, no differences in anti-spike or anti-RBD IgA antibody levels were observed at 2-6 weeks post-second dose (**Figure 8A, B**). However, by 6-months post-second dose, anti-spike and anti-RBD IgA antibody levels in HI vaccine recipients became higher than in uninfected participants (p < 0.0001 and p = 0.0011, respectively; **Figure 8A, B**). Anti-RBD IgA antibody levels in asymptomatic breakthrough recipients were likewise higher than those of uninfected participants (p = 0.037; **Figure 8B**).

### Assessment of Serological Neutralization of Live Ancestral SARS-CoV-2 over a Period of 6-Months Post-Second Dose

We lastly conducted live ancestral SARS-CoV-2 neutralization assays to characterize the neutralizing capacity of LTCH staff sera at 2-6 weeks and 6-months post-second dose. Mathematical models of exponential growth/decay were again fitted to individual values to evaluate response kinetics over time.

Across the entire cohort, LTCH staff sera had largely demonstrated robust neutralizing capacity against ancestral SARS-CoV-2 at 2-6 weeks post-second dose of BNT162b2 or mRNA-1273 (**Figure 9A**). However, with the wane of SARS-CoV-2 spike- and RBD-specific IgG and IgA antibody levels, serological neutralizing capacity likewise exhibited significant declines over the course of 6-months post-second dose (p < 0.0001; **Figure 9A**). As shown in **Figure 9D**, the rate of decay was observed to be bimodal; of the LTCH staff longitudinally analyzed, 99/113 (88%) experienced a relatively slower decay rate with a half-life that could not be reliably determined due to seemingly stationary response kinetics. Contrarily, the remaining 14/113 (12%) experienced a considerably faster median decay rate with a measurable half-life of 27 (26–28) days (**Table S9**).

**Figure 9:**
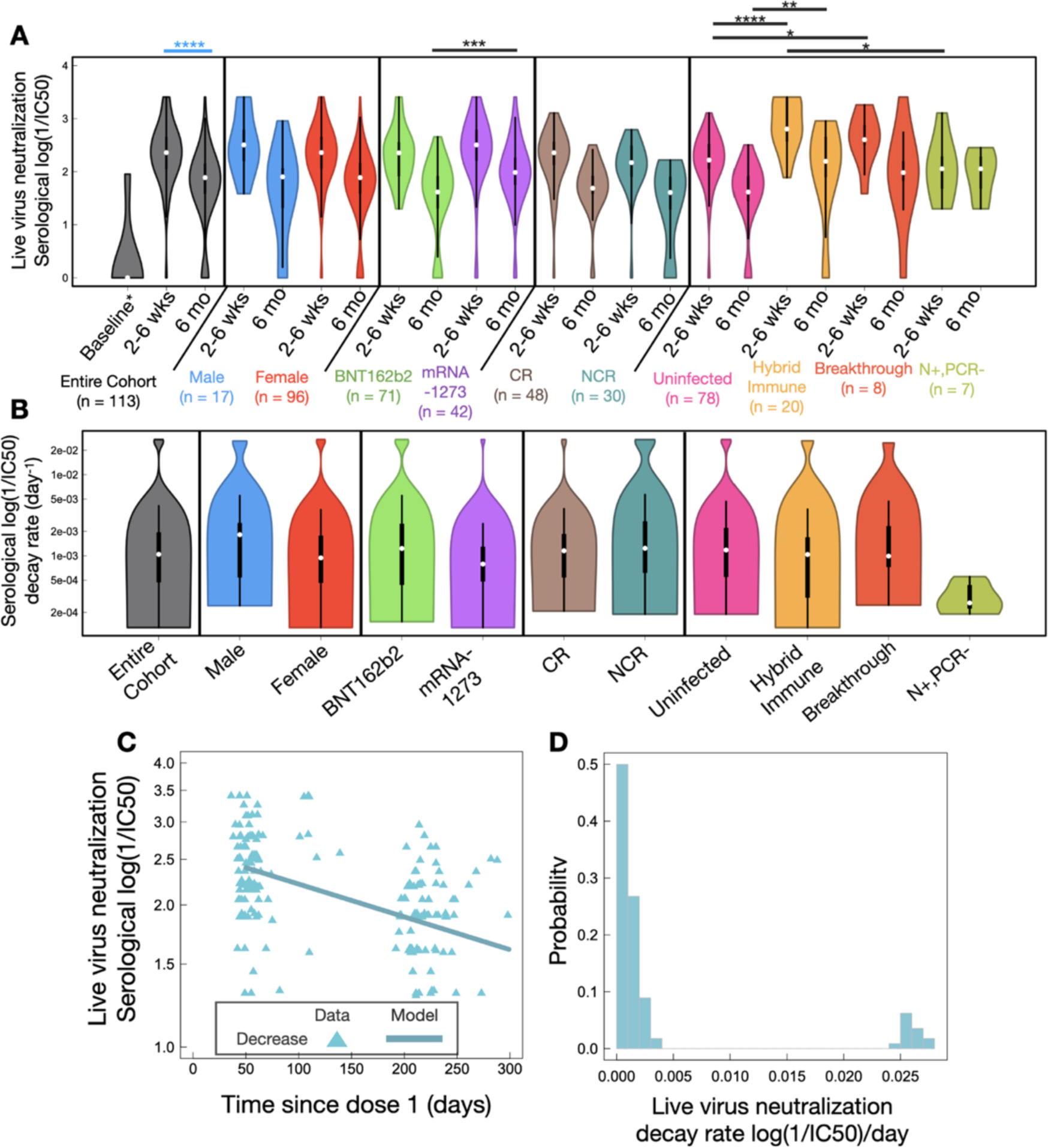
(A) violin plot analysis of serological neutralizing capacity against ancestral SARS-CoV-2 at 2-6 weeks and 6-months post-second dose. Squares and whiskers within each violin plot represent medians with IQRs. Refer to **Table S8** for medians with IQRs for each group. Significant cross-sectional comparisons are indicated with black asterisks. Significant longitudinal comparisons are displayed with blue asterisks for the entire cohort alone to avoid overcrowding the figure. Baseline is comprised of a group of 7 uninfected LTCH staff. (B) violin plot analysis of decay rates in serological neutralizing capacity across the entire LTCH staff cohort and following adjustments for sex, vaccine type, and immunological phenotype. (C) Mathematical modelling of decay in serological neutralizing capacity since dose 1 of BNT162b2 or mRNA-1273. (D) A histogram of individual decay rates in serological neutralizing capacity. Refer to **Table S9** for a summary of median decay rates with half-lives.

Stratifying for sex revealed no statistically significant difference in durability (**Figure 9B**; **Table S9**) or median serological neutralizing capacity (**Figure 9A**) between males and females.

Upon stratifying for vaccine type, the serological neutralization of live ancestral SARS-CoV-2 trended to be greater in recipients of mRNA-1273 than BNT162b2 at 2-6 weeks post-second dose, although statistical significance was not quite attained (p = 0.057; **Figure 9A**). However, despite similar decay kinetics between the two vaccine groups (**Figure 9B**; **Table S9**), serological neutralizing capacity later became significantly greater in recipients of mRNA-1273 (p = 0.0006) at 6-months post-second dose (**Figure 7A**).

Assessing the impacts of pre-existing T-cell cross-reactivity to SARS-CoV-2 on serological neutralizing capacity showed that CR vaccine recipients trended to exhibit greater neutralization than NCR recipients (p = 0.067) at 2-6 weeks post-second dose (**Figure 9A**). However, durability was not observed to differ between the two groups (**Figure 9B**; **Table S9**) and differences were entirely lost by 6-months post-second dose (**Figure9A**).

HI vaccinees exhibited superior serological neutralizing capacity compared to uninfected participants at both 2-6 weeks (p < 0.0001) and 6-months (p = 0.0038) post-second dose (**Figure 9A**). It was of additional interest to note that prior to the emergence of anti-N IgG or IgA antibodies, asymptomatic breakthrough vaccine recipients demonstrated significantly greater median neutralizing capacity against ancestral SARS-CoV-2 than uninfected participants at 2-6 weeks post-second dose (p = 0.049; **Figure 9A**).

The rate of decay in serological neutralizing capacity was observed to heavily overlap across all immunological phenotypes with the exception of N+, PCR(-) vaccine recipients, who interestingly demonstrated the slowest median decay rate (**Figure 9B**; **Table S9**). Due to seemingly stationary decay kinetics, half-lives could not be reliably computed for any immunological phenotype.

### Correlations between Spike-Specific T-cell and Humoral Responses to mRNA Vaccination with BNT162b2 or mRNA-1273

Having now characterized the cellular and humoral immune responses induced by mRNA vaccination, we next assessed how spike-specific IFN-γ and IL-2 T-cell responses in LTCH staff may have contributed to the magnitude and durability of vaccine-induced neutralizing antibody levels. To avoid potential confounding effects of natural infection, uninfected and hybrid immune participants were analyzed separately.

Notably, moderate to very strong positive correlations were observed between anti-spike and anti-RBD IgG/IgA antibody levels and serological neutralizing capacity. These relationships were generally strongest at 2-6 weeks post-second dose for HI vaccine recipients and at 6-months post-second dose for uninfected recipients (**Figure 10**). Accordingly, we first investigated how mRNA vaccine-induced spike-specific T-cell responses correlated with serological antibody levels at either timepoint. No correlations could be detected at either 2-6 weeks or 6-months post-second dose. However, within uninfected participants, initial spike-specific IL-2 T-cell responses at 2-6 weeks post-second dose were found to weakly correlate with serological anti-spike IgG (r_s_ = 0.3513; p = 0.0016), anti-RBD IgG (r_s_ = 0.2992; p = 0.0078), and anti-RBD IgA (r_s_ = 0.2412; p = 0.0334) antibody levels at the later 6-month timepoint (**Figure 11A**). Initial dual IFN-γ/IL-2 T-cell responses to spike at 2-6 weeks were likewise observed to weakly correlate with later anti-spike IgG antibody levels (r_s_ = 0.2594; p = 0.0218) at 6-months post-second dose (**Figure 11**). No such correlations could be observed for HI vaccine recipients (**Figure 11**).

**Figure 10:**
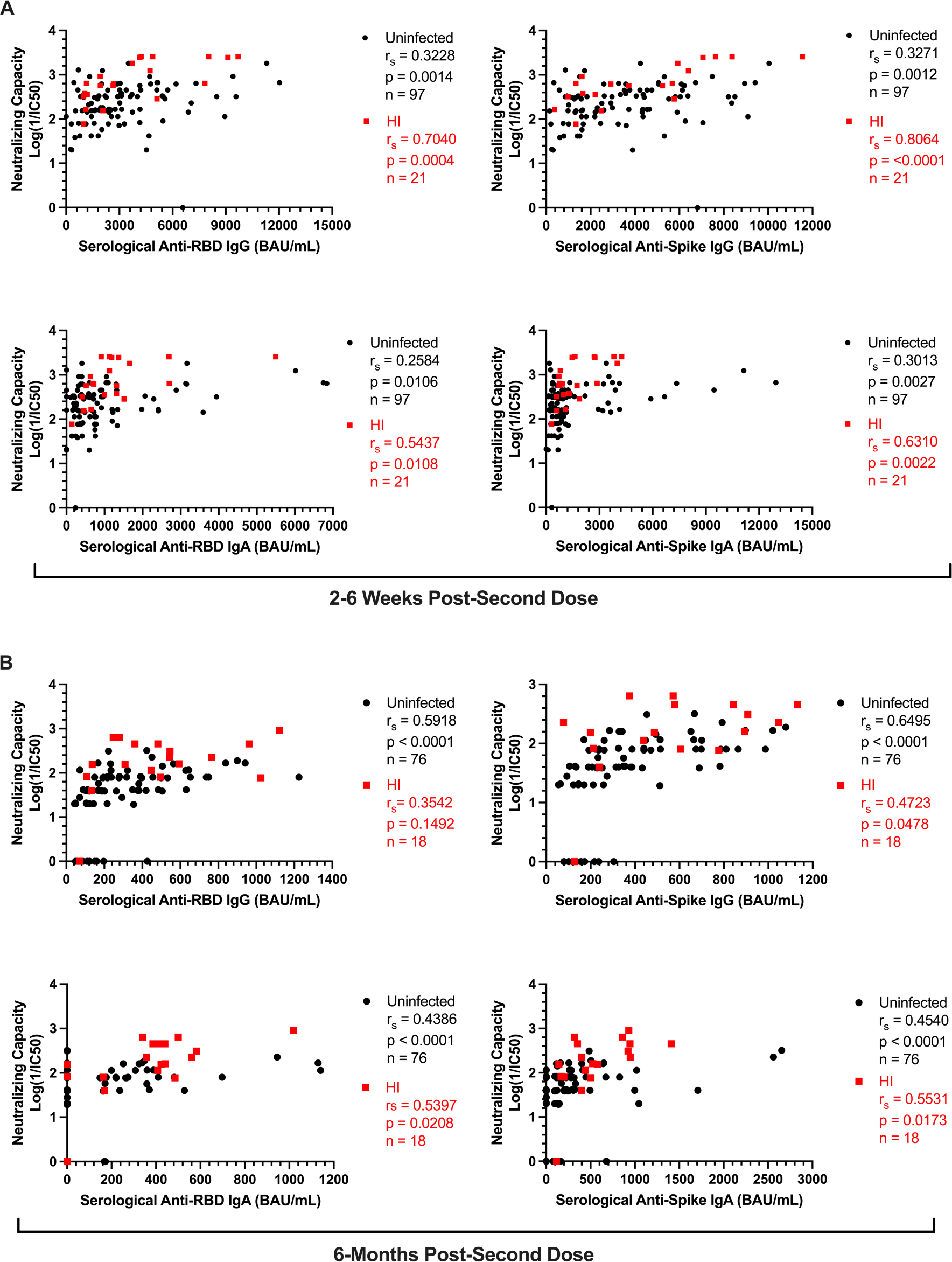
Correlations observed between serological anti-spike/RBD IgG and IgA antibody levels and neutralizing capacity against live ancestral SARS-CoV-2 in uninfected and HI vaccine recipients at (A) 2-6 weeks and (B) 6-months post-second dose of BNT162b2 or mRNA-1273. Black circles = uninfected; red squares = HI.

**Figure 11:**
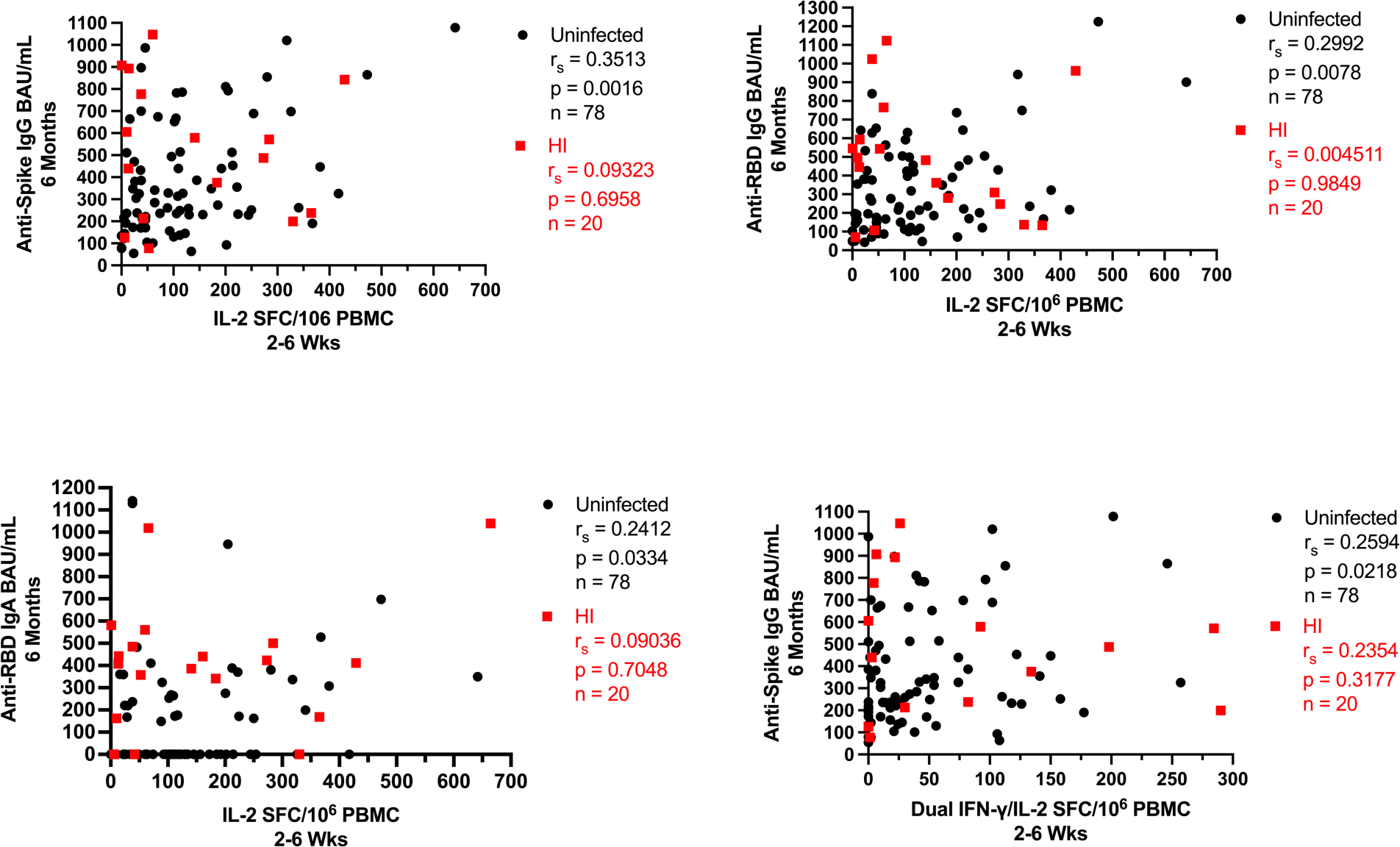
Spike-specific T-cell responses at 2-6 weeks post-second dose plotted against serological anti-spike IgG, anti-RBD IgG, and anti-RBD IgA antibody levels at 6-months post-second in uninfected and HI LTCH staff. Black circles = uninfected; red squares = HI.

We next investigated whether spike-specific T-cell responses directly correlated with serological neutralizing capacity. Within the HI vaccinee group, moderate positive correlations were observed between initial spike-specific IFN-γ (r_s_ = 0.4972, p = 0.0358) and dual IFN-γ/IL-2 (r_s_ = 0.5264, p = 0.0248) T-cell responses at 2-6 weeks post-second dose and serological neutralizing capacity at 6-months post-second dose (**Figure 12A**). Contrastingly, within uninfected vaccinees, only weak positive correlations could be detected between initial spike-specific IFN-γ (r_s_ = 0.2628; p = 0.0218), IL-2 (r_s_ = 0.2898; p = 0.0111), and dual IFN-γ/IL-2 (r_s_ = 0.3129; p = 0.0059) T-cell responses at 2-6 weeks post-second dose and serological neutralizing capacity at 6-months post-second dose (**Figure 12A**).

**Figure 12:**
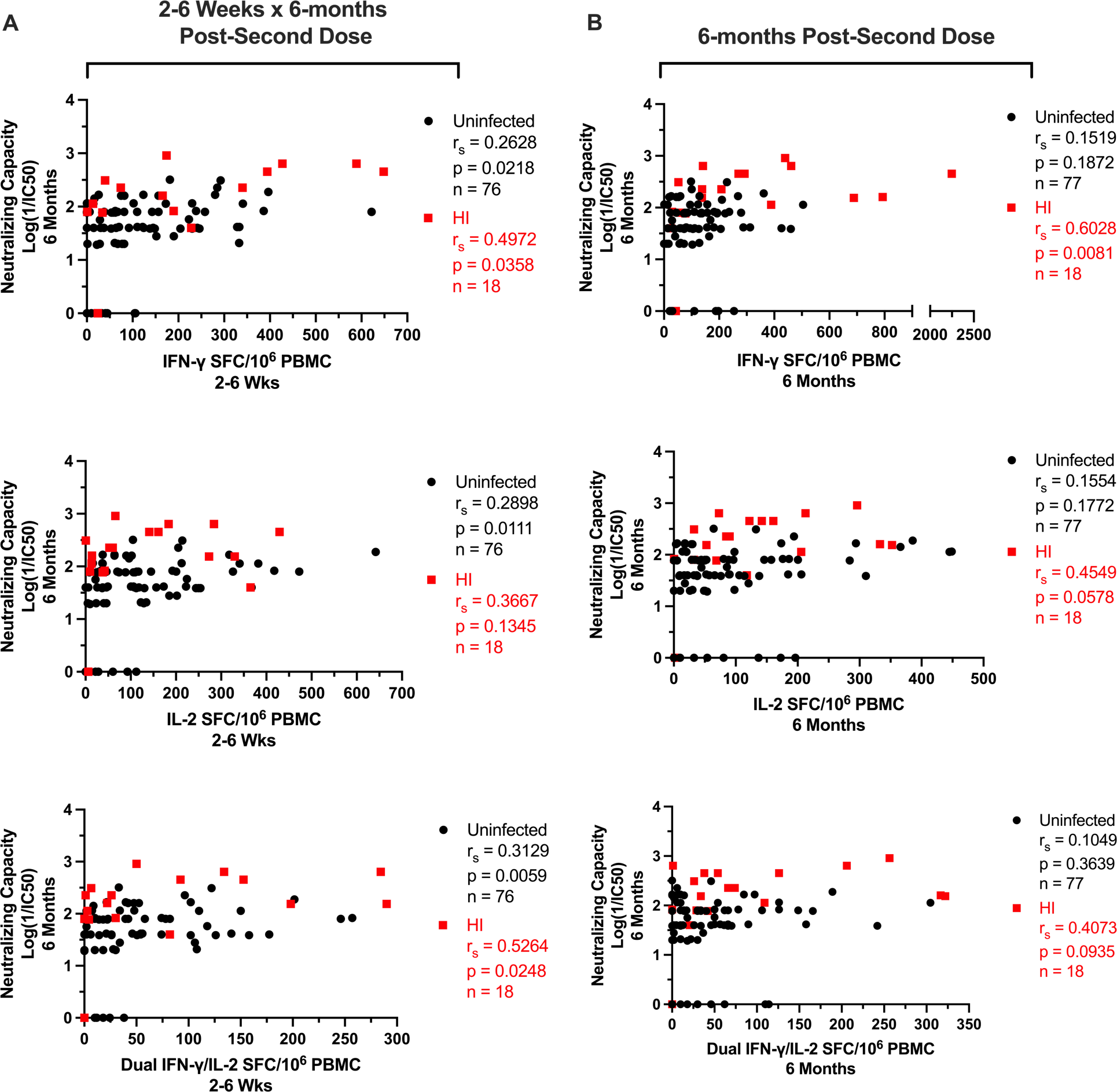
(A) Correlations between serological neutralizing capacity and net spike-specific IFN-γ, IL-2, and dual IFN-γ/IL-2 T-cell responses at 6-months post-second dose in uninfected and HI vaccine recipients. (B) Correlations between serological neutralizing capacity at 6-months post-second dose and net spike-specific IFN-γ, IL-2, and dual IFN-γ/IL-2 T-cell responses at 2-6 weeks post-second dose in uninfected and HI vaccine recipients. Black circles = uninfected; red squares = HI.

Moreover, at 6-months post-second dose, spike-specific IFN-γ T-cell responses were observed to strongly correlate (r_s_ = 0.6028, p = 0.0081) with serological neutralizing capacity in HI vaccine recipients (**Figure 12B**). Moderate positive correlations approaching significance were additionally observed for spike-specific IL-2 (r_s_ = 0.4549, p = 0.0578) and dual IFN-γ/IL-2 (r_s_ = 0.4073, p = 0.0935) T-cell responses (**Figure 12B**). However, no correlations could be observed for uninfected vaccine recipients at this timepoint (**Figure 12B**).

We lastly investigated whether initial spike-specific T-cell responses at 2-6 weeks post-second dose impacted the durability of serological neutralizing capacity. Interestingly, as shown in **Figure 13**, participants who experienced a slower rate of decay in serological neutralizing capacity (**Figure 9D**, n = 99) trended to exhibit greater median spike-specific IL-2 T-cell responses at 2-6 weeks post-second dose (p = 0.0749) than participants who experienced a faster rate of decay (**Figure 9D**, n = 14).

**Figure 13:**
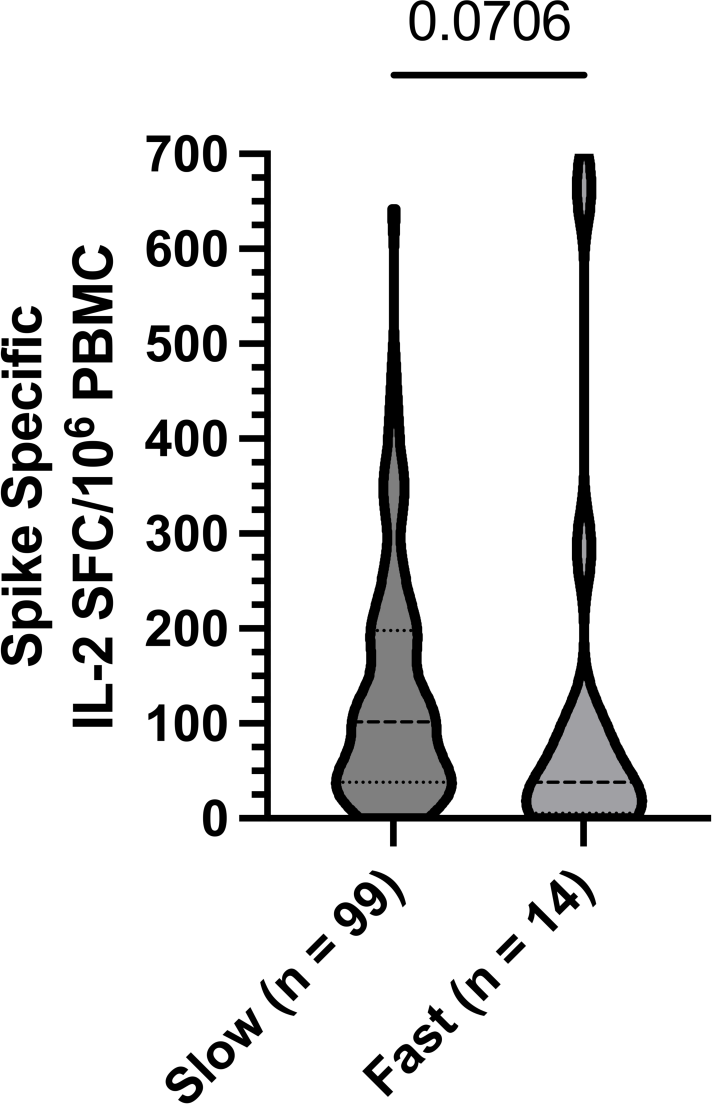
Comparisons of spike-specific (A) IL-2 T-cell responses and (B) IFN-γ/IL-2 T-cell response ratios at 2-6 weeks post-second dose between LTCH staff with slow and fast decay rates in serological neutralizing capacity. Broken lines represent medians with IQRs. Refer to **Table S9** for median decay rates with half-lives for each group.

Overall, mRNA vaccine-induced spike-specific IFN-γ, IL-2, and dual IFN-γ/IL-2 T-cell responses were observed to influence serological neutralizing capacity against ancestral SARS-CoV-2 within both uninfected and HI vaccine recipients over the course of 6-months post-second dose. Of the two cytokines assessed, IL-2 seemingly played a more pronounced role in promoting durability.

### Potential Off-target T-cell Responses Induced in Vaccine Recipients of BNT162b2 or mRNA-1273

Throughout the study, IFN-γ, IL-2, and dual IFN-γ/IL-2 non-spike-specific T-cell responses were continuously monitored by ELISpot in LTCH staff. In the following analyses, these non-spike-specific T-cell responses are reported as cumulative measures of nucleocapsid, envelope, membrane, and NSP-specific responses.

From 2-6 weeks to 6-months post-second dose, trending increases in non-spike-specific T-cell responses were noticed in the majority of vaccine recipients assessed (**Figure 14)**. Most notably, 21/30 (70%) NCR vaccine recipients began exhibiting non-spike-specific IFN-γ (p = 0.0004) and IL-2 (p = 0.0107) T-cell responses at 6-months post-second dose without ever reporting any clinical symptoms compatible with COVID-19, testing positive for COVID-19 by PCR/RAT, or seroconverting to possess anti-N IgG or IgA antibodies (**Figure 14**; **Figure S1**). This may suggest that a portion of NCR participants identified at 2-6 weeks post-second dose had possessed pre-existing cross-reactive T-cell memory to SARS-CoV-2 that was below detectable levels but later amplified by off-target vaccine effects.

**Figure 14:**
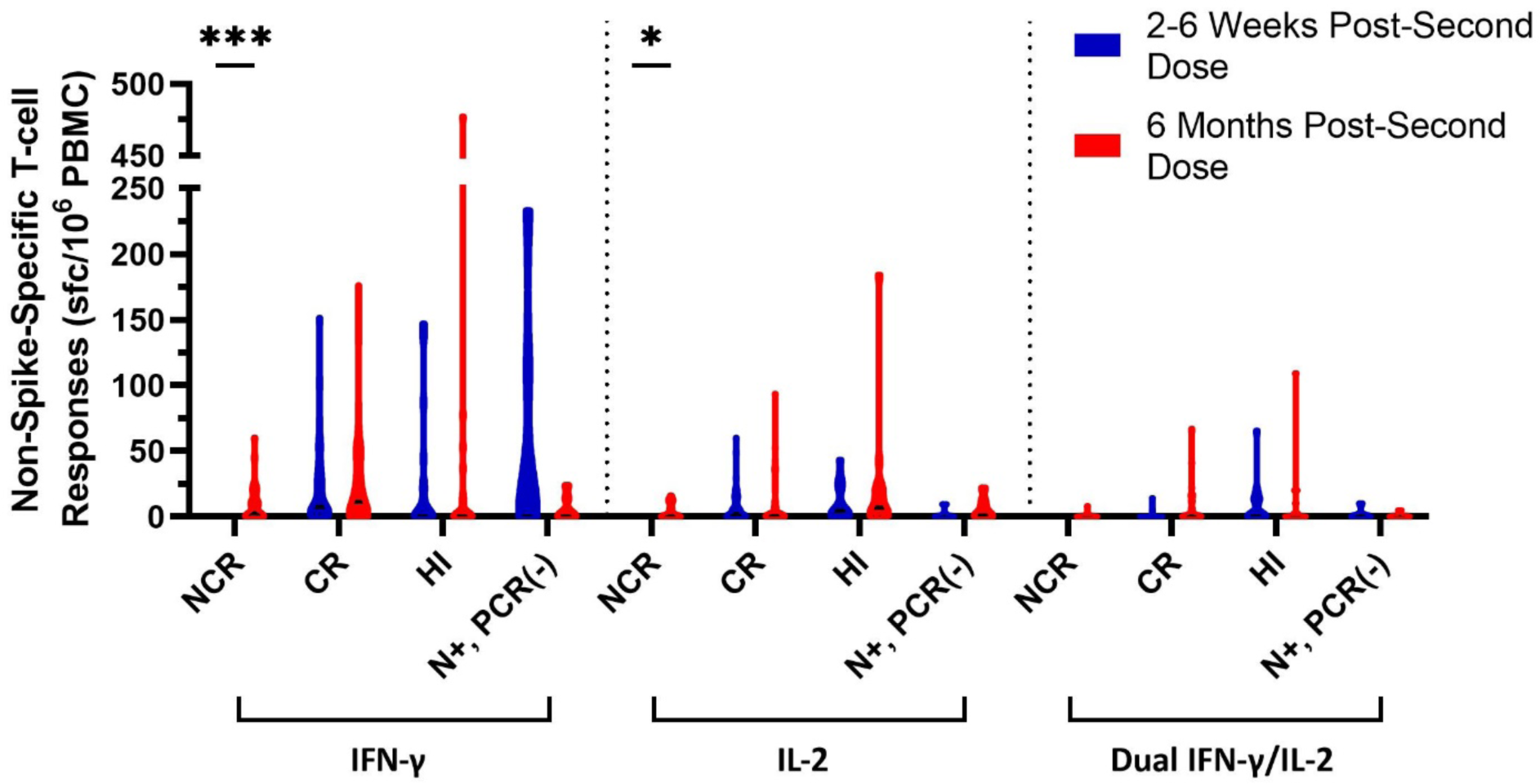
A longitudinal comparison of net non-spike-specific IFN-γ, IL-2, and dual IFN-γ/IL-2 T-cell responses from 2-6 weeks to 6-months post-second dose in NCR (n = 30), CR (n = 48), HI (n = 20), and N+, PCR(-) (n = 7) vaccine recipients. Non-spike-specific T-cell responses represent cumulative responses to nucleoprotein, envelope, membrane, and non-structural protein. Violin plots represent median responses with interquartile ranges.

To investigate this possibility, we first conducted longitudinal comparisons in CEF-, CEFTA-, and SEB-specific IFN-γ, IL-2, and dual IFN-γ/IL-2 T-cell responses from baseline to 6-months post-second dose. As shown in **Figure 15A**, baseline participants (n = 7) had generally exhibited significant increases in CEF- and CEFTA-specific IFN-γ, IL-2, and dual IFN-γ/IL-2 T-cell responses by 2-6 weeks post-second dose (p = 0.0461). However, no significant changes in SEB-specific T-cell responses could be observed (**Figure 15A**).

**Figure 15:**
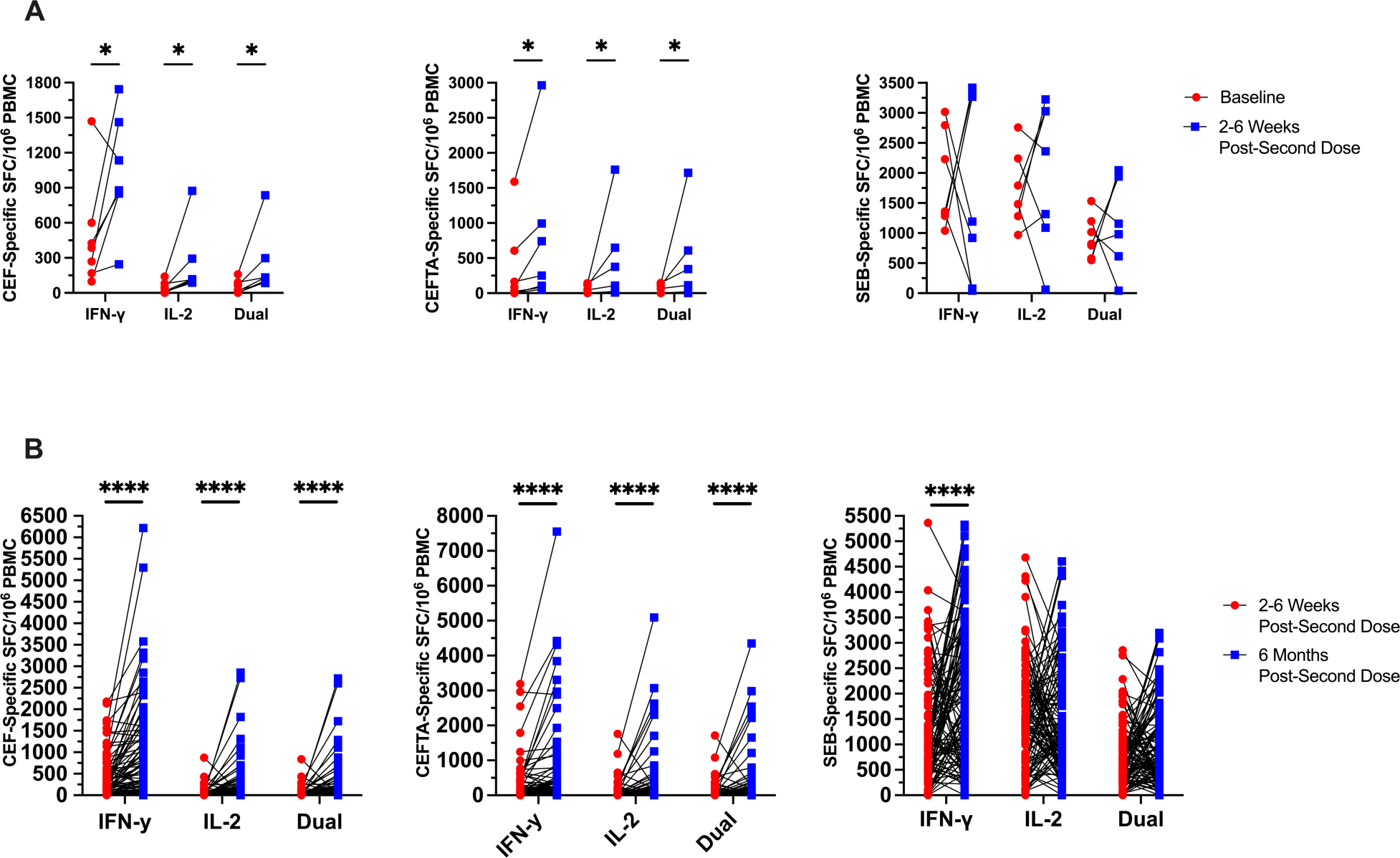
(A) Longitudinal pairwise comparisons (n = 7) of net CEF-, CEFTA-, and SEB-specific IFN-γ, IL-2, and dual IFN-γ/IL-2 T-cell responses from baseline (red circles) to 2-6 weeks post-second dose (blue squares). (B) Longitudinal pairwise comparisons (n = 105) of net CEF-, CEFTA-, and SEB-specific IFN-γ, IL-2, and dual IFN-γ/IL-2 T-cell responses from 2-6 weeks (red circles) to 6-months post-second dose (blue squares).

Increases in both CEF- and CEFTA-specific T-cell responses became most striking when comparisons were made between 2-6 weeks and 6-months post-second dose (p < 0.0001; **Figure 15B**). Highly significant increases were likewise observed for SEB-specific IFN-γ (p < 0.0001), but not IL-2 or dual IFN-γ/IL-2 T-cell responses (**Figure 15B**).

We next investigated whether undetectable non-spike-specific T-cell responses in NCR recipients at 2-6 weeks post-second dose could be amplified *in vitro*. For this experiment, 4 NCR recipients were randomly selected. Given that all participants in our study were previously exposed to HCoV (**Figure 3**), our highly conserved NSP masterpool (34) was used to screen for undetectable T-cell cross-reactivity at 2-6 weeks post-second dose. Briefly, PBMC samples were expanded for 7-days under stimulation with NSP, CEF, CEFTA, or SEB. T-cell responses were then assessed against these antigens by ELISpot (see methods). Although the 7-day *in vitro* expansion had substantially amplified CEF-, CEFTA-, and SEB-specific IFN-γ and IL-2 T-cell responses in all four NCR recipients, NSP-specific T-cell responses only became detectable in 1/4 (25%) recipients at 2-6 weeks post-second dose (data not shown).

## DISCUSSION

T-cells are key players in the adaptive immune response to invading pathogens and are essential for the development of neutralizing antibody titers (12, 15, 16). While several previous studies have evaluated the associations between SARS-CoV-2-specific CD4^+^ T-cell responses, serological antibody titers, and neutralization within the context of natural infection (4, 5, 20–22), very few studies to date have evaluated these associations within the context of vaccination (23, 24). The impact of pre-existing cross-reactive T-cell immunity to SARS-CoV-2 has additionally only been investigated in few studies (33) and its precise effects on the kinetics of cellular and humoral responses to vaccination are yet unclear. The aim of this study was to thoroughly investigate the associations between mRNA vaccine-induced spike-specific IFN-γ and IL-2 T-cell responses and neutralizing antibody development. The impacts of pre-existing cross-reactive T-cell immunity to SARS-CoV-2 on the magnitude and kinetics of cellular and humoral responses to vaccination were also investigated.

We began our investigation by first characterizing T-cell and humoral responses induced by mRNA vaccination with BNT162b2 or mRNA-1273 in our LTCH staff cohort. As expected, relative to baseline, robust spike-specific IFN-γ and IL-2 T-cell responses, anti-spike/RBD IgG and IgA antibody levels, and serological neutralization of ancestral SARS-CoV-2 were observed at 2-6 weeks post-second dose. In line with previous studies, although vaccine-induced humoral responses significantly waned over the course of 6-months post-second dose, spike-specific T-cell responses across the entire cohort were well preserved (58–65). Interestingly, mathematical modelling of T-cell response kinetics over the course of 6-months post-second dose revealed a gradual growth of spike-specific IFN-γ and IL-2 T-cell responses in 54% and 46% of vaccinees, respectively. A gradual decrease was contrastingly observed in all remaining vaccine recipients. To our knowledge, no previous studies have reported similar findings. However, a plausible explanation may be ‘cellular sensitization without seroconversion’, whereby SARS-CoV-2-specific T-cell responses are expanded in response to viral exposure in the absence of seroconversion, COVID-19-related symptoms, or positive PCR testing (66–68). The likelihood of this phenomenon for our LTCH staff cohort is increased by the fact that our cohort worked and lived in high-risk conditions during the COVID-19 pandemic.

Mathematical modelling additionally revealed bimodal decay kinetics in the serological neutralization of ancestral SARS-CoV-2 across the entire LTCH staff cohort – while the decay in neutralizing antibody levels was highly gradual in 88% of individuals, the remaining 12% experienced considerably faster decay rates with a median half-life of 27 (26–28) days (**Figure 9D**). As shown in **Figure 13**, participants who experienced slower decay rates in neutralization trended to exhibit greater spike-specific IL-2 T-cell responses at 2-6 weeks post-second dose than those with faster decay rates. Accordingly, initial spike-specific IL-2 T-cell responses to vaccination were potentially predictive of the durability of serological neutralizing capacity. Previous work has similarly shown a significant positive correlation between mRNA vaccine-induced spike-specific IL-2 T-cell responses 2-4 weeks post-second dose and neutralizing antibody levels 3-4 months post-second dose in recipients with immune-mediated inflammatory diseases and in healthy controls (69).

We had additionally asserted direct positive correlations between initial spike-specific T-cell responses at 2-6 weeks post-second dose and serological neutralizing capacity at 6-months post-second dose that were considerably strengthened in HI vaccine recipients (**Figure 12**). Spike-specific T-cell responses trended to be greatest in HI recipients at 2-6 weeks post-second dose (**Figure 5**). Hybrid immunity may have thus boosted durability in a subset of participants and contributed to the bimodal decay kinetics across the entire LTCH staff cohort. Previous studies have similarly reported enhanced durability in neutralizing antibody levels in hybrid immune vaccine recipients after a 2-dose mRNA vaccine regimen (70, 71). Anti-N+, PCR(-) participants interestingly exhibited the greatest durability in serological neutralizing capacity, with no measurable decay or growth over time. Based on a previous report, it is possible that these vaccine recipients had contracted recent SARS-CoV-2 infections that elevated their neutralizing antibody levels at baseline but impeded their humoral responses to the second dose of BNT162b2 or mRNA-1273 (72). Given that no COVID-19-related symptoms or positive PCR tests were previously reported in anti-N+, PCR(-) recipients, this would imply that asymptomatic infections may be capable of blunting the humoral immune response to mRNA vaccination. Further studies would be needed to assess the impacts of asymptomatic SARS-CoV-2 infections on humoral immune responses to mRNA vaccines.

We next investigated how pre-existing T-cell cross-reactivity to SARS-CoV-2 may have impacted the magnitude and kinetics of T-cell and humoral responses induced by mRNA vaccination. Within our study, CR vaccinees were a sub-stratification of uninfected participants who exhibited significant spike-specific T-cell responses by ELISpot to any SARS-CoV-2 masterpool at baseline or to any non-spike masterpool at 2-6 weeks post-second dose. We understood that cellular sensitization without seroconversion may potentially explain the non-spike-specific T-cell responses exhibited by CR vaccine recipients. A comparison between HI and CR vaccinees showed that CR recipients predominantly mounted non-spike-specific T-cell responses against the NSP masterpool at 6-months post-second dose. These observations strongly aligned with Swadling et al. who reported a preferential expansion of T-cell responses against NSP7 (contained within our NSP masterpool) and NSP12 – highly conserved antigenic targets across human common-cold coronaviruses – in exposed seronegative healthcare workers who repeatedly tested negative for COVID-19 by PCR (66). Akin to their study, we additionally observed a preferential expansion of T-cell responses toward the membrane masterpool in HI vaccine recipients, which is contrastingly considerably less conserved across human common-cold coronaviruses (66). Accordingly, with regards to non-spike-specific T-cell responses after vaccination, CR vaccinees were phenotypically distinct from HI recipients.

CR recipients exhibited significant boosts in spike-specific IFN-γ and dual IFN-γ/IL-2 T-cell responses at 2-6 weeks post-second dose compared to NCR vaccinees. Yet, consistent with a previous study (38), the boosts were transient and seemingly dissipated by 6-months post-second dose. In agreement with (39), we also could not detect significant differences in anti-spike/RBD IgG antibody levels between CR and NCR vaccine recipients. Although serological neutralizing capacity trended to be greater in CR vaccinees at 2-6 weeks post-second dose, differences were lost by 6-months post-second dose. This contrasted with (39), which continuously detected elevated serological neutralizing capacity in cross-reactive vaccine recipients by 6-months post-second dose. Discrepancies may be explained by the administration of a lower 25ug dose of mRNA-1273 in (39) compared to the standard 100ug dose administered in our study. A higher 100ug of mRNA-1273 may have elicited a more potent response that ‘drowned out’ the effects of pre-existing cross-reactivity on neutralizing capacity. Our pooling of BNT162b2 and mRNA-1273 participants in our analysis may have also diminished our ability to detect differences between NCR and CR vaccinees. Nonetheless, pre-existing T-cell cross-reactivity to SARS-CoV-2 was not observed to impact the kinetics of mRNA vaccine-induced spike-specific IgG responses or serological neutralizing capacity.

Interestingly, anti-RBD IgA levels were transiently lower in CR than in NCR vaccine recipients at 2-6 weeks post-second dose. This may be attributable to the greater spike-specific IFN-γ T-cell responses observed in CR vaccinees at this timepoint – recent literature suggests significant associations between heightened type I and type II interferon signaling and delayed anti-RBD antibody kinetics in COVID-19 patients (73). CR vaccine recipients were additionally the only immunological phenotype to have experienced significant declines in spike-specific IL-2 T-cell responses from 2-6 weeks to 6-months post-second dose. Whether this is due to the emigration of cells from the circulation to specific tissues (74, 75) requires further study.

From 2-6 weeks to 6-months post-second dose, most vaccine recipients exhibited trending increases in non-spike-specific T-cell responses to SARS-CoV-2. These increases were especially prominent and had reached statistical significance within NCR vaccine recipients. Within NCR recipients, we hypothesized that the emergence of non-spike-specific T-cell responses to SARS-CoV-2 at 6-months post-second dose was attributable to the vaccine-induced amplification of previously undetectable responses. Of four NCR vaccinees tested at 2-6 weeks post-second dose, T-cell responses to our conserved NSP peptide masterpool could only be amplified to detectable levels in one recipient. However, given that all recipients in our study exhibited evidence of exposure to previous common-cold coronaviruses (**Figure 3**), it is still likely that the remaining 3/4 NCR recipients tested possessed cross-reactive T-cell memory to SARS-CoV-2 that was not captured using our NSP peptide masterpool. Interestingly, highly significant increases in CEF-, CEFTA-, and SEB-specific T-cell responses were additionally observed across the entire LTCH staff cohort by 6-months post-second dose. This may potentially be due to bystander activation, which can reportedly induce immune responses to non-related antigens following vaccination or natural infection (76–78).

In summary, our data highlights the potential contributions of mRNA vaccine-induced SARS-CoV-2 spike-specific T-cell responses to serological neutralizing capacity within both uninfected and HI vaccine recipients up to 6-months post-second dose. IL-2 was observed to play a particularly pronounced role in promoting the durability of neutralizing antibodies. Pre-existing cross-reactive T-cell immunity to SARS-CoV-2 had only a modest effect on the magnitude and kinetics of T-cell and humoral responses to mRNA vaccination. These findings should ultimately inform the development of novel pan-coronavirus vaccines that can minimize the impacts of both current and future human coronaviruses of concern.

## ACKNOWLEDGEMENTS

This work was supported by funding from the Canadian Institute of Health Research, the COVID-19 Immunity Task Force, and NRC’s Pandemic Response Challenge Program.

Sample acquisition, coordination, and transportation were performed by Anjali Patel, Keelia Quinn de Launay, Jamie Boyd, and Alison Takaoka under the direction of Sharon Straus at Unity Health, St. Michael’s Hospital, Toronto.

PBMC and serum sample preparation was conducted with the assistance of Justin Mendoza, Serena Chau, Thomas Morningstar, Erik Mihelic, and Joseph Vincent De Fazio at the Ostrowski Lab, University of Toronto.

Chemiluminescent ELISA assays for SARS-CoV-2-specific IgG and IgA antibody detection in serum were conducted by the Network Biology Collaborative Centre at Lunenfeld-Tanenbaum Research Institute of Sinai Health, Toronto (Adrian Pasculescu, Freda Qi, Melanie Delgado-Brand, Tulunay R. Tursun, Geneviève Mailhot, Roya Monica Dayam, and Karen Colwill), a facility supported by the Canada Foundation for Innovation, the Ontario Government, Genome Canada and Ontario Genomics (OGI-139).

Chemiluminescent ELISAs assays for HCoV spike-specific IgG antibody detection in serum were conducted by the University of Ottawa Serology and Diagnostics High Throughput Facility (Danielle Dewar-Darch, Gwendoline Ward, Justino Hernadez-Soto, Abishek Xavier, Nicholas Bradette, Klaudia Baumann).

Live SARS-CoV-2 neutralization assays were conducted by Patrick Budylowski and Serena Chau from the University of Toronto.

Mathematical modelling and statistics were conducted with the aid of Chapin S. Korosec and Jane M. Heffernen at the Modelling Infection and Immunity Lab and the Center for Disease Modelling, Mathematics and Statistics Department, York University.

A special thanks to Mario Ostrowski for providing the resources, direction, and expertise necessary to make much of this work possible. Additional thanks to Feng Yun Yue for the management of laboratory operations.

## SUPPLEMENTARY MATERIALS

**Table S1:**
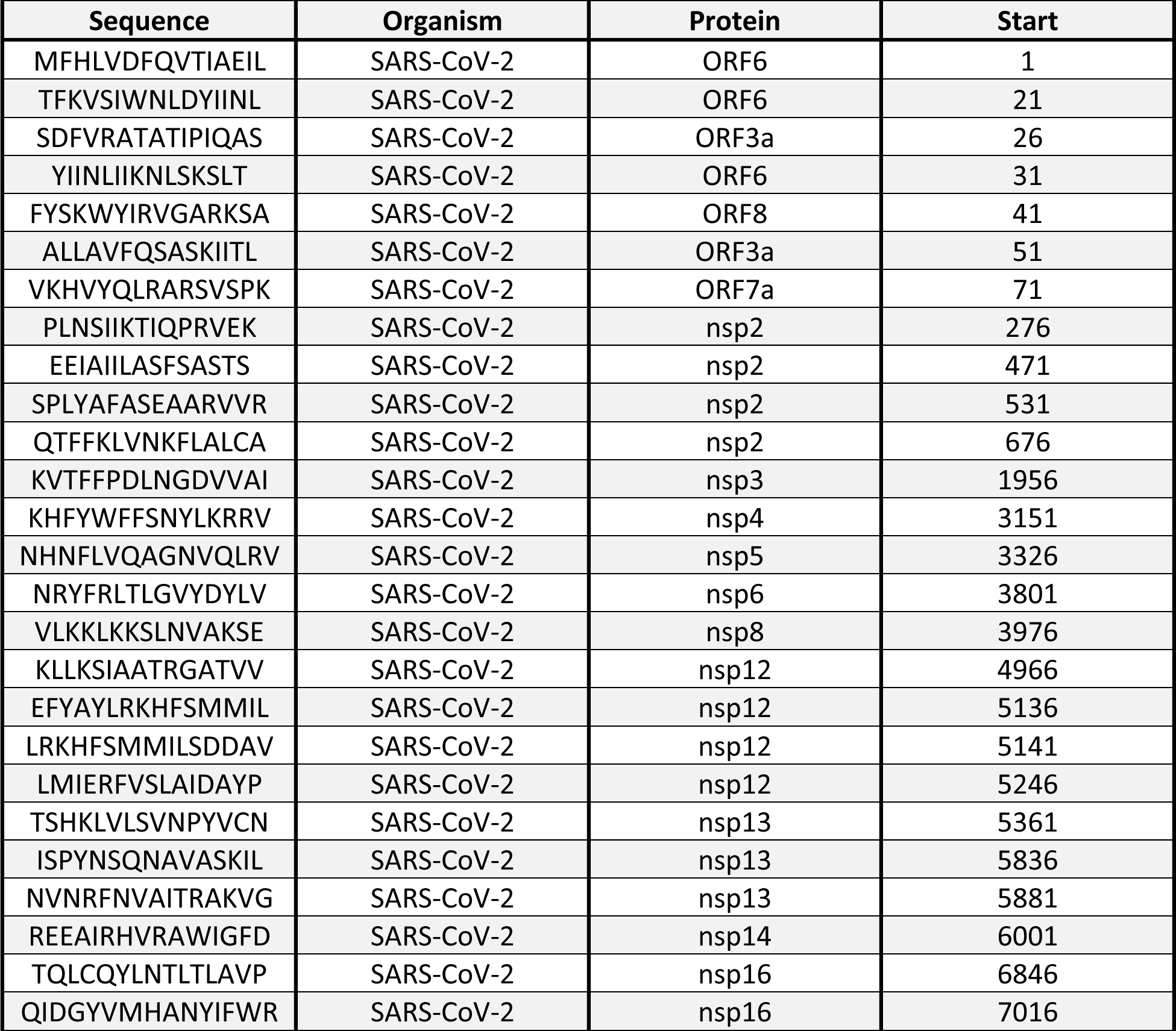
15-mer peptides constituting the NSP masterpool.

**Table S2:** Clinical characteristics of hybrid immune LTCH staff analyzed.

| Participant ID | # of PCR-Confirmed Infections | Date of Most Recent Positive PCR Test (yyyy-mm-dd) | Date of 1<sup>st</sup> Vaccine Dose | Date of 2<sup>nd</sup> Vaccine Dose | Days between Infection and 1<sup>st</sup> Dose | Days between Infection and 2<sup>nd</sup> Dose | Severity of Infection<sup>1</sup> | Date of Blood Draw 2-6 Weeks Post-Second Dose (yyyy-mm-dd) |
| --- | --- | --- | --- | --- | --- | --- | --- | --- |
| RSS002 | 1 | 2020-04-07 | 2021-01-03 | 2021-01-31 | -271 | -299 | 2 | 2021-02-20 |
| CVS001 | 1 | 2020-04-25 | 2021-01-06 | 2021-01-27 | -256 | -277 | 2 | 2021-02-11 |
| CCS004 | 1 | 2020-05-22 | 2021-01-19 | 2021-02-16 | -242 | -270 | 2 | 2021-03-10 |
| CWS001 | 1 | 2020-09-25 | 2021-03-17 | 2021-02-25 | -173 | -153 | 2 | 2021-03-18 |
| VSS009 | 1 | 2020-10-01 | 2021-02-25 | 2021-05-19 | -147 | -230 | 1 | 2021-06-15 |
| DLS019 | 1 | 2020-11-02 | 2021-03-25 | 2021-06-03 | -143 | -213 | 1 | 2021-06-23 |
| VSS007 | 1 | 2020-10-13 | 2021-01-18 | 2021-02-15 | -97 | -125 | 1 | 2021-03-11 |
| VSS004 | 1 | 2020-10-01 | 2021-01-05 | 2021-02-02 | -96 | -124 | 1 | 2021-02-20 |
| VSS003 | 1 | 2020-10-18 | 2021-01-10 | 2021-02-15 | -84 | -120 | 2 | 2021-03-02 |
| VSS008 | 1 | 2020-10-16 | 2021-01-05 | 2021-02-02 | -81 | -109 | 2 | 2021-03-02 |
| ONS010 | 1 | 2021-01-11 | 2021-03-15 | 2021-06-04 | -63 | -144 | 2 | 2021-06-28 |
| NLS010 | 1 | 2021-01-10 | 2021-02-21 | 2021-05-22 | -42 | -132 | 2 | 2021-06-10 |
| CVS002 | 1 | 2020-11-25 | 2021-01-04 | 2021-02-01 | -40 | -68 | 1 | 2021-02-18 |
| NLS003 | 1 | 2021-01-13 | 2021-02-20 | 2021-05-21 | -38 | -128 | 2 | 2021-06-10 |
| CVS004 | 1 | 2021-01-04 | 2021-01-04 | 2021-02-01 | 0 | -28 | 2 | 2021-02-18 |
| DLS018 | 1 | 2021-01-06 | 2021-01-04 | 2021-01-25 | 2 | -19 | 0 | 2021-03-02 |
| CVS003 | 1 | 2021-01-17 | 2021-01-13 | 2021-02-17 | 4 | -31 | 1 | 2021-03-11 |
| CCS006 | 1 | 2021-02-10 | 2021-01-19 | 2021-02-26 | 22 | -16 | 1 | 2021-03-18 |
| MCS035 | 1 | 2021-01-20 | 2020-12-28 | 2021-02-05 | 23 | -16 | 2 | 2021-03-04 |
| DLS013<sup>2</sup> | 1 | 2021-02-09 | 2021-01-05 | 2021-01-26 | 35 | 14 | 0 | 2021-02-20 |

<sup>1</sup> 0 = Asymptomatic; 1 = Mild symptoms, no fever; 2 = Mild symptoms with fever, not hospitalized
<sup>2</sup> Exhibited an asymptomatic PCR-confirmed breakthrough infection 14 days following second vaccine dose, but 11 days prior to sample acquisition

**Table S3:**
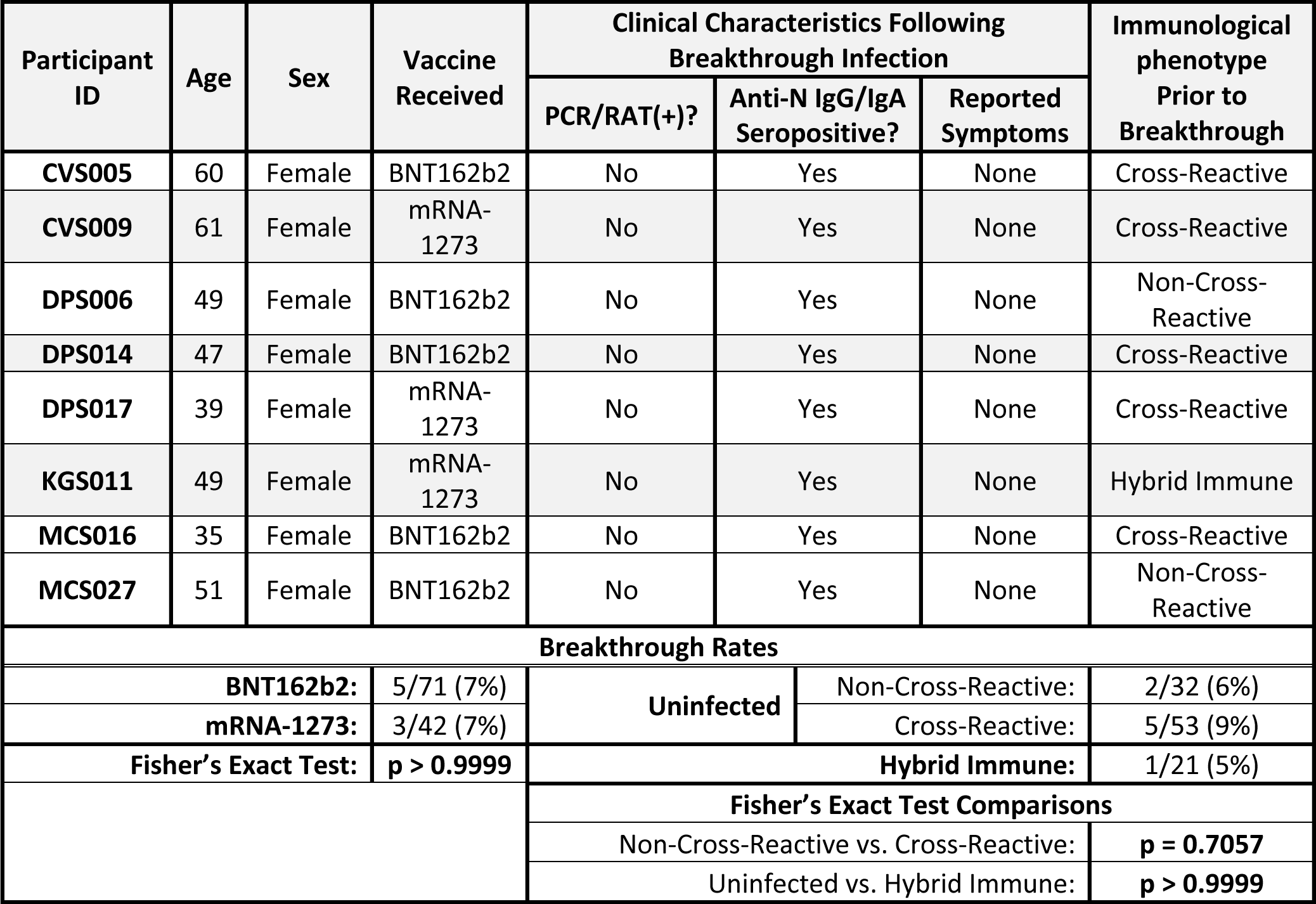
A summary of clinical characteristics of breakthrough vaccinees following infection.

**Table S4:** Spike-specific IFN-γ, IL-2, and dual IFN-γ/IL-2 T-cell responses following mRNA vaccination at 2-6 weeks and 6-months post-second dose.

| | Cytokine | Median Net Spike-Specific sfc/$10^6$ PBMC (IQR) | | | | | | | | | |
| --- | --- | --- | --- | --- | --- | --- | --- | --- | --- | --- | --- |
|  |  | Entire Cohort | Sex |  | Vaccine Type |  | Immunological phenotype |  |  |  | N+ PCR(-) (n = 7) |
|  |  |  | Male (n = 17) | Female (n = 96) | BNT162b2 (n = 71) | mRNA-1273 (n = 42) | Uninfected |  | HI (n = 20) | Breakthrough (n = 8) |  |
|  |  |  |  |  |  |  | CR (n = 48) | NCR (n = 30) |  |  |  |
| Baseline (n = 7) | IFN-γ | 0 (2.88) | - | - | - | - | - | - | - | - | - |
|  | IL-2 | 0 (0.172) | - | - | - | - | - | - | - | - | - |
| | Dual IFN-γ/IL-2 | 0 (0.00) | - | - | - | - | - | - | - | - | - |
| 2-6 Weeks Post 2<sup>nd</sup> Dose (n = 113) | IFN-γ | 106 (192.81) | 151.74 (279.89) | 95 (195.60) | 82 (154.03) | 141.35 (260.6) | 131.17 (178.32) | 59.09 (65.83) | 181.86 (417.07) | 129.35 (167.13) | 118 (186.24) |
|  | IL-2 | 96.34 (154.44) | 122.16 (223.66) | 94.52 (140.72) | 62.94 (102.35) | 158.66 (212.43) | 107.17 (162.99) | 76.16 (95.88) | 61 (174.21) | 109.52 (122.29) | 84.1 (140.7) |
| | Dual IFN-γ/IL-2 | 34.1 (74.56) | 38 (64.51) | 31.52 (76) | 25 (49.72) | 52 (103.06) | 38.89 (71.44) | 14.98 (43) | 44 (157.93) | 47.35 (29) | 58 (86.53) |
| 6-Months Post 2<sup>nd</sup> Dose (n = 113) | IFN-γ | 98 (140) | 128.69 (115.31) | 92.17 (143.37) | 85.37 (120.98) | 125.72 (200.69) | 107.52 (120.42) | 80.86 (115.50) | 191.17 (380.47) | 70.20 (125.47) | 63.08 (45.72) |
|  | IL-2 | 64.34 (109.31) | 73.03 (140.35) | 62.69 (105.32) | 50 (99.35) | 96 (159.60) | 51.17 (94.09) | 84.34 (138.67) | 106 (159.77) | 55.52 (48.50) | 49.37 (47) |
| | Dual IFN-γ/IL-2 | 30 (56) | 36.69 (120.97) | 30 (52) | 30 (54.49) | 34 (73.91) | 30 (45.92) | 26 (57) | 48 (121.33) | 18 (59.74) | 30 (18.80) |

**Table S5:** A summary of median growth and decay rates in spike-specific T-cell responses in LTCH staff from 2-6 weeks to 6-months post-second dose of BNT162b2 or mRNA-1273.

| | | IFN-γ | | | IL-2 | | |
| --- | --- | --- | --- | --- | --- | --- | --- |
|  |  | Entire Cohort<br>(n = 113) | BNT162b2<br>(n = 71) | mRNA-1273<br>(n = 42) | Entire Cohort<br>(n = 113) | BNT162b2<br>(n = 71) | mRNA-1273<br>(n = 42) |
| Growth | Median Growth (d<sup>-1</sup>) | 0.0038 | 0.0039 | 0.0038 | 0.0036 | 0.0039 | 0.0031 |
|  | Growth IQR (d<sup>-1</sup>) | 0.0007 | 0.0007 | 0.0008 | 0.0021 | 0.0023 | 0.0018 |
|  | Doubling Time (d) | 181.4583 | 179.7446 | 181.4583 | 192.9480 | 179.8388 | 224.6161 |
|  | Doubling Time<br>Range Max | 222.7788 | 218.8063 | 230.1476 | 471.7893 | 441.5696 | 525.4339 |
|  | Doubling Time<br>Range Min | 153.0677 | 152.5169 | 149.7728 | 121.2724 | 112.9124 | 142.8389 |
| Decay | Median Decay (d<sup>-1</sup>) | 0.0065 | 0.0057 | 0.0074 | 0.0058 | 0.0062 | 0.0051 |
|  | Decay IQR (d<sup>-1</sup>) | 0.0079 | 0.0066 | 0.0083 | 0.0061 | 0.0071 | 0.0044 |
|  | Half-Life (d) | 0.0057 | 0.0054 | 0.0061 | 0.0048 | 0.0051 | 0.0042 |
|  | Half-Life Range Max | 106.6929 | 121.3064 | 94.1327 | 118.9570 | 111.9496 | 135.1262 |
|  | Half-Life Range Min | 831.3968 | 2019.7067 | 565.6474 | 666.6370 | 616.9858 | 709.8587 |

**Table S6:** Serological anti-spike IgG, anti-RBD IgG, anti-spike IgA, and anti-RBD IgA antibody levels following mRNA vaccination at 2-6 weeks and 6-months post-second dose.

|  | Serum Antibody Titer | Median Binding Antibody Units/mL (IQR) |  |  |  |  |  |  |  |  |  |
| --- | --- | --- | --- | --- | --- | --- | --- | --- | --- | --- | --- |
|  |  | Entire Cohort | Sex |  | Vaccine Type |  | Immunological phenotype |  |  |  |  |
|  |  |  | Male (n = 17) | Female (n = 96) | BNT162b2 (n = 71) | mRNA-1273 (n = 42) | Uninfected |  | HI (n = 20) | Breakthrough (n = 8) | N+ PCR(-) (n = 7) |
|  |  |  |  |  |  |  | CR (n = 48) | NCR (n = 30) |  |  |  |
| Baseline (n = 7) | Anti-Spike IgG | 0 (0) | - | - | - | - | - | - | - | - | - |
|  | Anti-RBD IgG | 0 (0) | - | - | - | - | - | - | - | - | - |
|  | Anti-Spike IgA | 0 (0) | - | - | - | - | - | - | - | - | - |
|  | Anti-RBD IgA | 0 (0) | - | - | - | - | - | - | - | - | - |
| 2-6 Weeks Post 2<sup>nd</sup> Dose (n = 113) | Anti-Spike IgG | 2906.67 (3781.65) | 3889.54 (4199.9) | 2906.67 (3745.84) | 2411.78 (3721.21) | 4047.36 (4070.78) | 2782.39 (4083.33) | 2450.12 (2450.70) | 4480.59 (5561.67) | 4307.73 (2821.82) | 2411.78 (2020.76) |
|  | Anti-RBD IgG | 2554.69 (3114.49) | 3389.74 (3307.7) | 2547.97 (2792.07) | 2192.15 (2997.17) | 3072.60 (2912.36) | 2660.19 (3604.1) | 2391.16 (1639.18) | 2658.85 (3588.56) | 3295.86 (2931.96) | 2541.24 (935.79) |
|  | Anti-Spike IgA | 779.27 (1185.64) | 874.77 (572.72) | 741.83 (1226.67) | 589.73 (716.87) | 1187.46 (2368.06) | 592.03 (729.88) | 827.02 (1603.41) | 1193.39 (1964.25) | 764.31 (1187.97) | 1379.3 (1429.67) |
|  | Anti-RBD IgA | 648.84 (862.39) | 913.5 (618.42) | 614.6 (862.64) | 415.1 (497.16) | 1164.72 (912.50) | 426.04 (665.48) | 730.71 (822.63) | 1062.45 (686.79) | 660.76 (911.28) | 1094.16 (1294.33) |
| 6-Months Post 2<sup>nd</sup> Dose (n = 113) | Anti-Spike IgG | 341.56 (454.29) | 273.61 (433.73) | 348.06 (451.77) | 261.13 (262.27) | 552.07 (527.37) | 320 (260.76) | 261.23 (332.40) | 592.44 (601.40) | 451.93 (550.91) | 287.04 (322.05) |
|  | Anti-RBD IgG | 293.05 (399.50) | 235.33 (394.39) | 313.59 (393.84) | 216.62 (310.60) | 468.69 (410.91) | 270.27 (310.56) | 224.96 (336.64) | 489.36 (542.97) | 439.49 (595.11) | 449.19 (471.56) |
|  | Anti-Spike IgA | 249.74 (376.54) | 240.4 (169.36) | 250.26 (383.76) | 191.06 (299.57) | 323.94 (472.58) | 192.87 (220.24) | 216.06 (209.90) | 513.95 (532.68) | 288.33 (476.15) | 296.19 (503.67) |
|  | **Anti-RBD**<br>**IgA** | 148.34<br>(360.57) | 0<br>(388.87) | 155.33<br>(358.11) | 0<br>(234.87) | 327.70<br>(379.72) | 0<br>(224.89) | 0<br>(271.95) | 417.43<br>(217.33) | 224.90<br>(353.68) | 0<br>(595.53) |

**Table S7:**
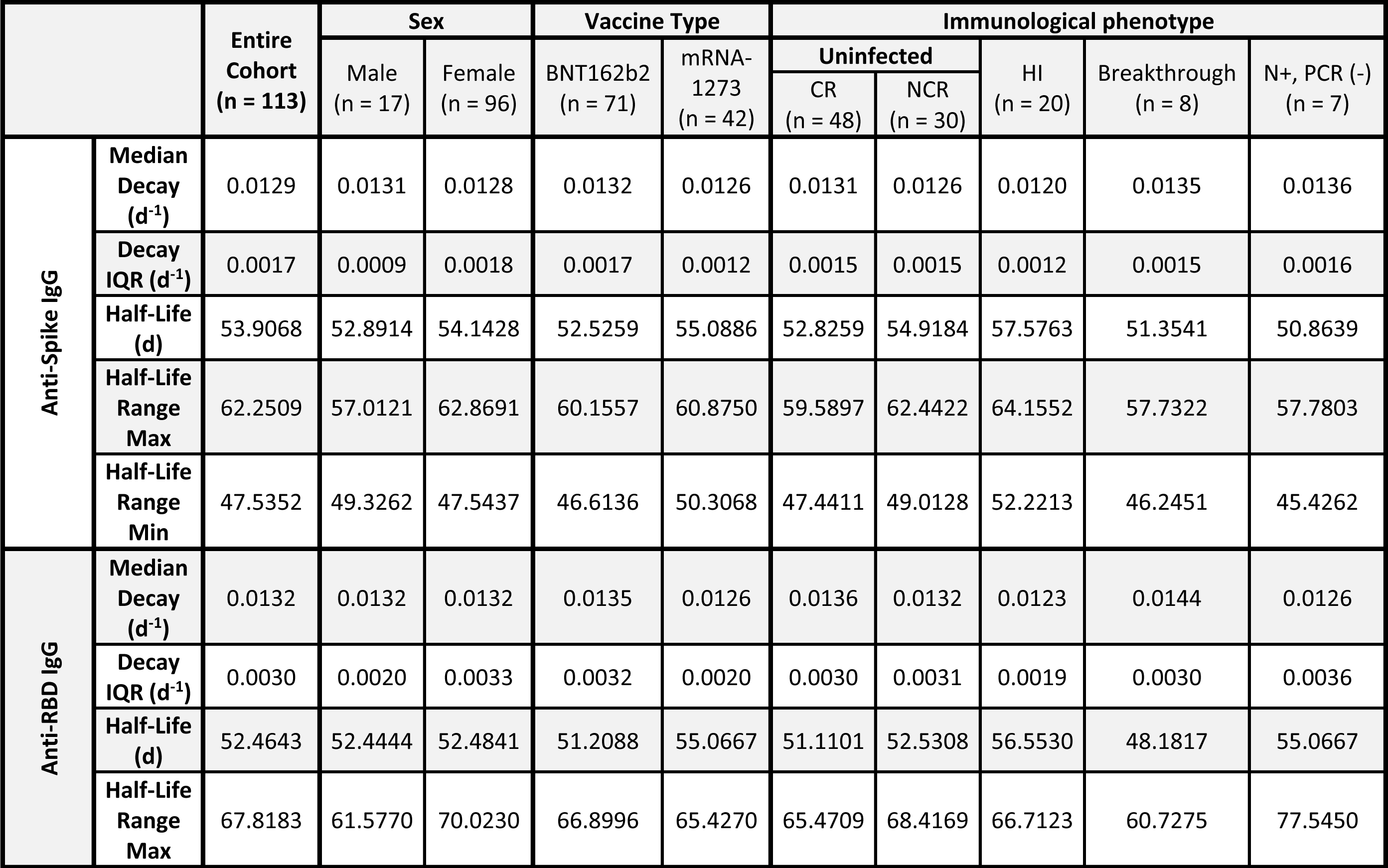
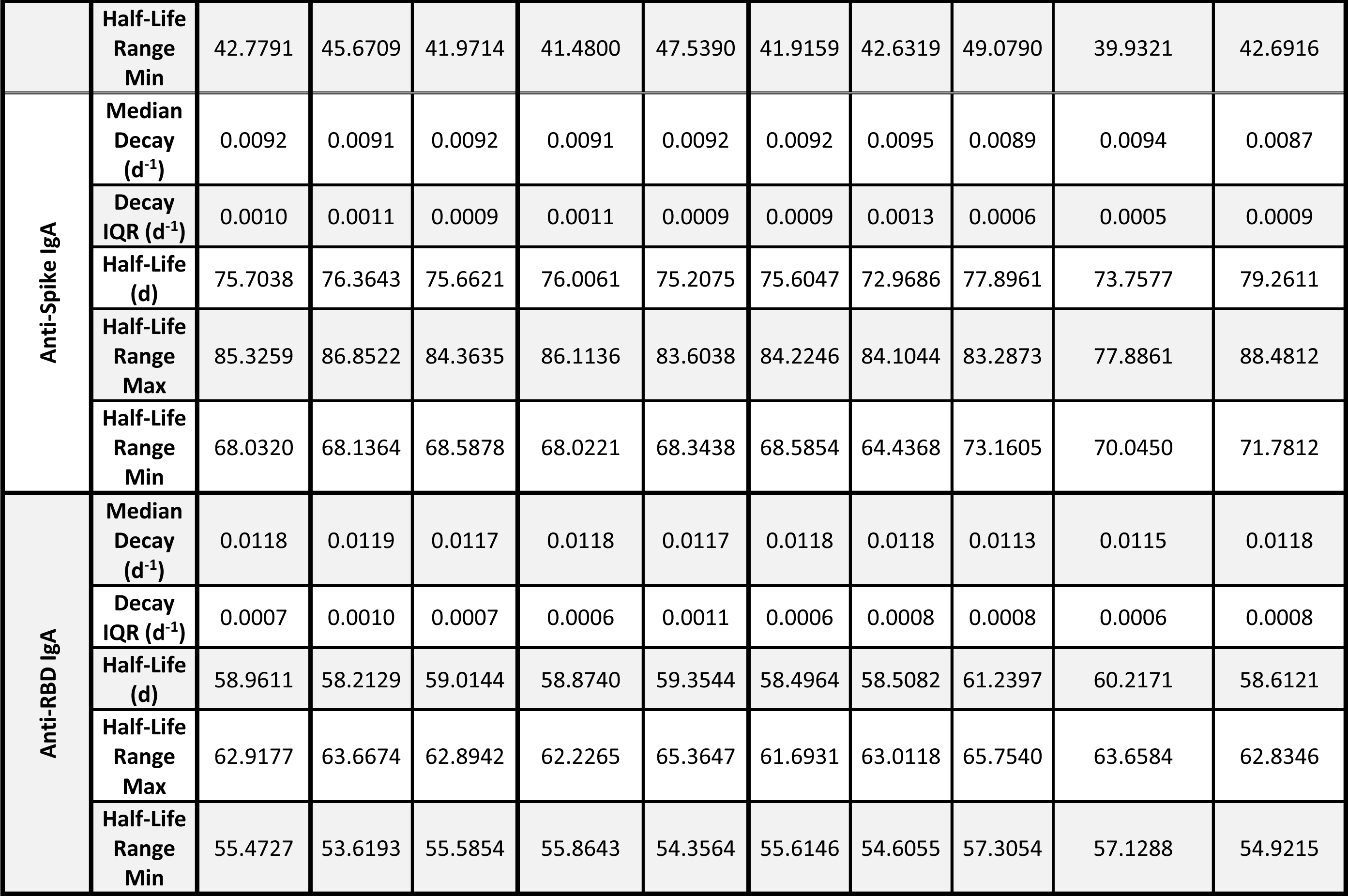
A summary of median decay rates in serological anti-spike and anti-RBD IgG and IgA antibody levels in vaccinated LTCH staff from 2-6 weeks to 6-months post-second dose of BNT162b2 or mRNA-1273.

**Table S8:** Serological neutralization of live ancestral SARS-CoV-2 *in vitro* following mRNA vaccination at 2-6 weeks and 6-months post-second dose of BNT162b2 or mRNA-1273.

|  | Median Log(1/IC50) (IQR) |  |  |  |  |  |  |  |  |  |
| --- | --- | --- | --- | --- | --- | --- | --- | --- | --- | --- |
|  | Entire Cohort | Sex |  | Vaccine Type |  | Immunological phenotype |  |  |  |  |
|  |  | Male<br>(n = 17) | Female<br>(n = 96) | BNT162b2<br>(n = 71) | mRNA-1273<br>(n = 42) | Uninfected |  | HI<br>(n = 20) | Breakthrough<br>(n = 8) | N+ PCR(-)<br>(n = 7) |
|  |  |  |  |  |  | CR<br>(n = 48) | NCR<br>(n = 30) |  |  |  |
| Baseline<br>(n = 7) | 0<br>(0) | - | - | - | - | - | - | - | - | - |
| 2-6 Weeks Post 2<sup>nd</sup> Dose<br>(n = 113) | 2.36<br>(0.60) | 2.51<br>(0.59) | 2.36<br>(0.60) | 2.35<br>(0.64) | 2.51<br>(0.58) | 2.36<br>(0.43) | 2.17<br>(0.59) | 2.81<br>(0.82) | 2.60<br>(0.37) | 2.05<br>(0.6) |
| 6-months Post 2<sup>nd</sup> Dose<br>(n = 113) | 1.89<br>(0.57) | 1.90<br>(0.75) | 1.89<br>(0.58) | 1.62<br>(0.61) | 1.99<br>(0.51) | 1.69<br>(0.33) | 1.61<br>(0.61) | 2.20<br>(0.76) | 1.98<br>(0.37) | 2.05<br>(0.6) |

**Table S9:** A summary of decay rates in serological neutralizing capacity against ancestral SARS-CoV-2 in vaccinated LTCH staff from 2-6 weeks to 6-months post-second dose of BNT162b2 or mRNA-1273.

|  |  | Entire Cohort<br>(n = 113) | Sex |  | Vaccine Type |  | Immunological phenotype |  |  |  |  |
| --- | --- | --- | --- | --- | --- | --- | --- | --- | --- | --- | --- |
|  |  |  | Male<br>(n = 17) | Female<br>(n = 96) | BNT162b2<br>(n = 71) | mRNA-1273<br>(n = 42) | Uninfected |  | HI<br>(n = 20) | Breakthrough<br>(n = 8) | N+ PCR(-)<br>(n = 7) |
|  |  |  |  |  |  |  | CR<br>(n = 48) | NCR<br>(n = 30) |  |  |  |
| Consolidated | Median Decay (d<sup>-1</sup>) | - | 0.0018 | 0.0009 | 0.0012 | 0.0008 | 0.0012 | 0.0012 | 0.0010 | 0.0010 | 0.0003 |
|  | Decay IQR (d<sup>-1</sup>) | - | 0.0020 | 0.0013 | 0.0020 | 0.0008 | 0.0013 | 0.0020 | 0.0014 | 0.0016 | 0.0002 |
| Slow (n = 99) | Median Decay (d<sup>-1</sup>) | 0.0008 | - | - | - | - | - | - | - | - | - |
|  | Decay IQR (d<sup>-1</sup>) | 0.0011 | - | - | - | - | - | - | - | - | - |
| Fast (n = 14) | Median Decay (d<sup>-1</sup>) | 0.0253 | - | - | - | - | - | - | - | - | - |
|  | Decay IQR (d<sup>-1</sup>) | 0.0012 | - | - | - | - | - | - | - | - | - |
|  | Half-Life (d) | 27.3941 | - | - | - | - | - | - | - | - | - |
|  | Half-Life Range Min | 26.1362 | - | - | - | - | - | - | - | - | - |
|  | Half-Life Range Max | 28.7793 | - | - | - | - | - | - | - | - | - |

**Figure S1:**
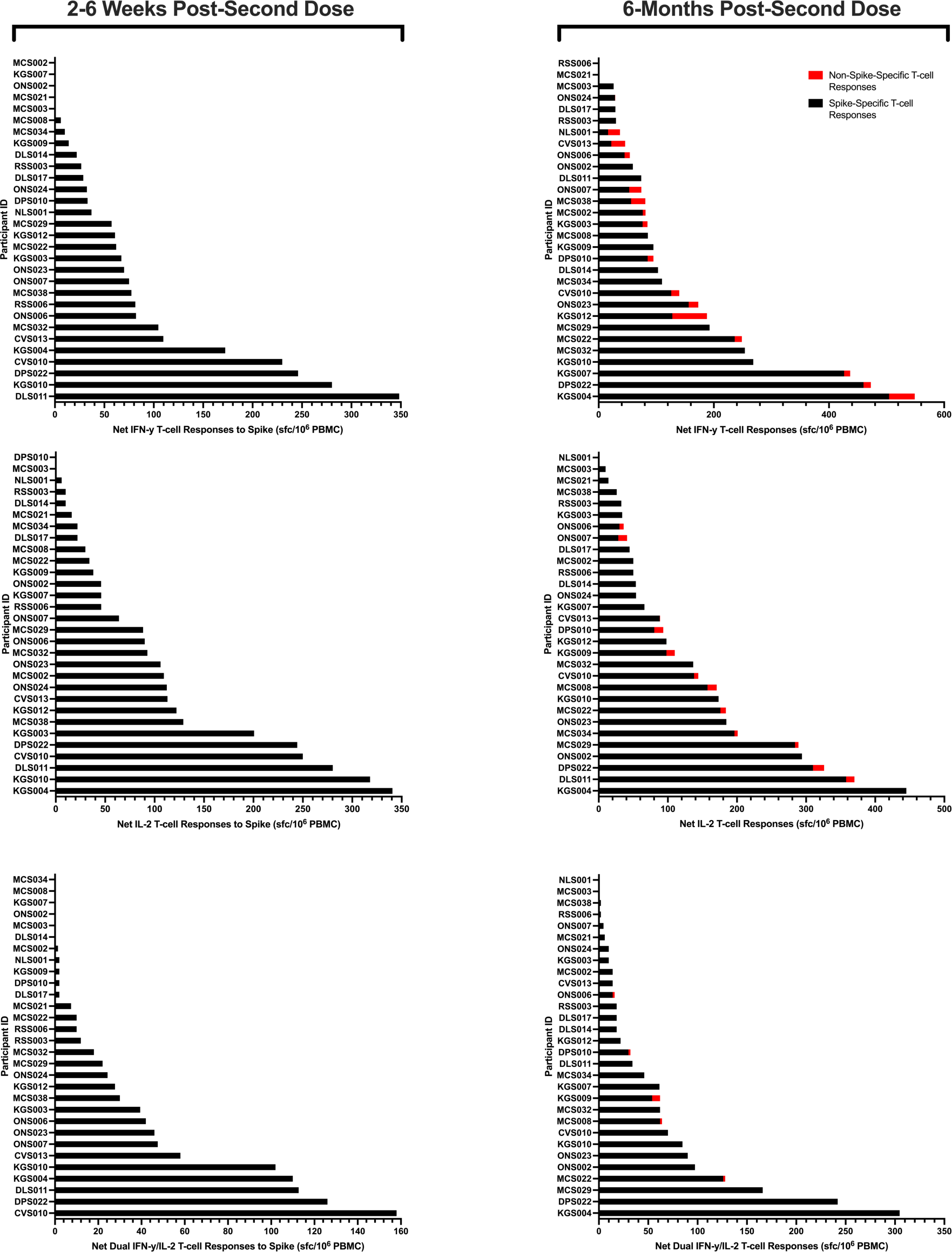
A summary of spike-specific (black) and non-spike-specific (red) T-cell responses in NCR vaccinees at 2-6 weeks and 6-months post-second dose. Non-spike-specific T-cell responses are cumulative of N-, E-, M-, and NSP-specific responses.

## REFERENCES

1. Goldblatt D, Alter G, Crotty S, Plotkin SA. 2022. Correlates of protection against SARS-CoV-2 infection and COVID-19 disease. Immunol Rev 310:6–26.

2. Murphy K, Weaver C, Berg L. 2022. Janeway’s Immunobiology10th Edition. W. W. Norton & Company, Inc., New York.

3. Tarke A, Potesta M, Varchetta S, Fenoglio D, Iannetta M, Sarmati L, Mele D, Dentone C, Bassetti M, Montesano C, Mondelli MU, Filaci G, Grifoni A, Sette A. 2022. Early and Polyantigenic CD4 T Cell Responses Correlate with Mild Disease in Acute COVID-19 Donors. Int J Mol Sci 23:7155.

4. Nelson RW, Chen Y, Venezia OL, Majerus RM, Shin DS, Carrington MN, Yu XG, Wesemann DR, Moon JJ, Luster AD, Abayneh BA, Allen P, Alter G, Antille D, Armstrong K, Balazs A, Bals J, Barbash M, Bartsch Y, Boucau J, Boyce S, Braley J, Branch K, Broderick K, Carney J, Chan A, Chevalier J, Chowdhury F, Daley G, Davidson S, Dougan M, Drew D, Einkauf K, Elliman A, Fallon J, Fedirko L, Finn K, Flaherty K, Flannery J, Forde P, Garcia-Broncano P, Gettings E, Golan D, Griffin A, Grimmel S, Grinke K, Hall K, Hartana C, Healy M, Heller H, Henault D, Holland G, Jiang C, Jilg N, Kaplonek P, Karpell M, Kayitesi C, Lam EC, LaValle V, Lefteri K, Lian X, Lichterfeld M, Lingwood D, Liu H, Liu J, Lu Y, Luthern S, Ly N, Marchewka J, Martino B, McNamara R, Michell A, Millstrom I, Miranda N, Nambu C, Nelson S, Noone M, O’Callaghan C, Ommerborn C, Osborn M, Pacheco LC, Phan N, Pillai S, Porto FA, Rassadkina Y, Reissis A, Rosenthal A, Ruzicka F, Ryan E, Seiger K, Selleck K, Sessa L, Sharpe A, Sharr C, Shin S, Singh N, Slaughenhaupt S, Sheppard KS, Sun W, Sun X, Suschana E, Ticheli H, Trocha-Piechocka A, Wilson V, Wong C, Worrall D, Zhu A, Manickas-Hill Z, DeMers E, Judge K, Ghebremichael MS, Lai P, Li J, Yu XG, Walker B, Carrington M, Martin M, Yuki Y. 2022. SARS-CoV-2 epitope–specific CD4+ memory T cell responses across COVID-19 disease severity and antibody durability. Sci Immunol 7:eabl9464.

5. Rydyznski Moderbacher C, Ramirez SI, Dan JM, Grifoni A, Hastie KM, Weiskopf D, Belanger S, Abbott RK, Kim C, Choi J, Kato Y, Crotty EG, Kim C, Rawlings SA, Mateus J, Tse LPV, Frazier A, Baric R, Peters B, Greenbaum J, Ollmann Saphire E, Smith DM, Sette A, Crotty S. 2020. Antigen-Specific Adaptive Immunity to SARS-CoV-2 in Acute COVID-19 and Associations with Age and Disease Severity. Cell 183:996–1912.E19.

6. Lin J, Law R, Korosec CS, Zhou C, Koh WH, Ghaemi MS, Samaan P, Ooi HK, Matveev V, Yue F, Gingras A-C, Estacio A, Buchholz M, Cheatley P Lou, Mohammadi A, Kaul R, Pavinski K, Mubareka S, McGeer AJ, Leis JA, Heffernan JM, Ostrowski M. 2022. Longitudinal Assessment of SARS-CoV-2-Specific T Cell Cytokine-Producing Responses for 1 Year Reveals Persistence of Multicytokine Proliferative Responses, with Greater Immunity Associated with Disease Severity. J Virol 96:e0050922.

7. Zeng B, Gao L, Zhou Q, Yu K, Sun F. 2022. Effectiveness of COVID-19 vaccines against SARS-CoV-2 variants of concern: a systematic review and meta-analysis. BMC Med 20:200.

8. Kurhade C, Zou J, Xia H, Liu M, Chang HC, Ren P, Xie X, Shi PY. 2023. Low neutralization of SARS-CoV-2 Omicron BA.2.75.2, BQ.1.1 and XBB.1 by parental mRNA vaccine or a BA.5 bivalent booster. Nat Med 29:344–347.

9. Tseng HF, Ackerson BK, Luo Y, Sy LS, Talarico CA, Tian Y, Bruxvoort KJ, Tubert JE, Florea A, Ku JH, Lee GS, Choi SK, Takhar HS, Aragones M, Qian L. 2022. Effectiveness of mRNA-1273 against SARS-CoV-2 Omicron and Delta variants. Nat Med 28:1063–1071.

10. Keeton R, Tincho MB, Ngomti A, Baguma R, Benede N, Suzuki A, Khan K, Cele S, Bernstein M, Karim F, Madzorera S V., Moyo-Gwete T, Mennen M, Skelem S, Adriaanse M, Mutithu D, Aremu O, Stek C, du Bruyn E, Van Der Mescht MA, de Beer Z, de Villiers TR, Bodenstein A, van den Berg G, Mendes A, Strydom A, Venter M, Giandhari J, Naidoo Y, Pillay S, Tegally H, Grifoni A, Weiskopf D, Sette A, Wilkinson RJ, de Oliveira T, Bekker LG, Gray G, Ueckermann V, Rossouw T, Boswell MT, Bihman J, Moore PL, Sigal A, Ntusi NAB, Burgers WA, Riou C. 2022. T cell responses to SARS-CoV-2 spike cross-recognize Omicron. Nature 603:488-492.

11. Mazzoni A, Vanni A, Spinicci M, Capone M, Lamacchia G, Salvati L, Coppi M, Antonelli A, Carnasciali A, Farahvachi P, Giovacchini N, Aiezza N, Malentacchi F, Zammarchi L, Liotta F, Rossolini GM, Bartoloni A, Cosmi L, Maggi L, Annunziato F. 2022. SARS-CoV-2 Spike-Specific CD4+ T Cell Response Is Conserved Against Variants of Concern, Including Omicron. Front Immunol 13:801431.

12. Kervevan J, Chakrabarti LA. 2021. Role of CD4+ T Cells in the Control of Viral Infections: Recent Advances and Open Questions. Int J Mol Sci 22:523.

13. Tang P, Hasan MR, Chemaitelly H, Yassine HM, Benslimane FM, Al Khatib HA, AlMukdad S, Coyle P, Ayoub HH, Al Kanaani Z, Al Kuwari E, Jeremijenko A, Kaleeckal AH, Latif AN, Shaik RM, Abdul Rahim HF, Nasrallah GK, Al Kuwari MG, Al Romaihi HE, Butt AA, Al-Thani MH, Al Khal A, Bertollini R, Abu-Raddad LJ. 2021. BNT162b2 and mRNA-1273 COVID-19 vaccine effectiveness against the SARS-CoV-2 Delta variant in Qatar. Nat Med 27:2136–2143.

14. McMenamin ME, Nealon J, Lin Y, Wong JY, Cheung JK, Lau EHY, Wu P, Leung GM, Cowling BJ. 2022. Vaccine effectiveness of one, two, and three doses of BNT162b2 and CoronaVac against COVID-19 in Hong Kong: a population-based observational study. Lancet Infect Dis 22:1435–1443.

15. Sette A, Crotty S. 2021. Adaptive immunity to SARS-CoV-2 and COVID-19. Cell 184:861–880.

16. Crotty S. 2019. T Follicular Helper Cell Biology: A Decade of Discovery and Diseases. Immunity 10.1016/j.immuni.2019.04.011.

17. Lucas C, Klein J, Sundaram ME, Liu F, Wong P, Silva J, Mao T, Oh JE, Mohanty S, Huang J, Tokuyama M, Lu P, Venkataraman A, Park A, Israelow B, Vogels CBF, Muenker MC, Chang C-H, Casanovas-Massana A, Moore AJ, Zell J, Fournier JB, Obaid A, Robertson AJ, Lu-Culligan A, Zhao A, Nelson A, Brito A, Nunez A, Martin A, Watkins AE, Geng B, Chun CJ, Kalinich CC, Harden CA, Todeasa C, Jensen C, Dorgay CE, Kim D, McDonald D, Shepard D, Courchaine E, White EB, Song E, Silva E, Kudo E, DeIuliis G, Rahming H, Park H-J, Matos I, Ott I, Nouws J, Valdez J, Fauver J, Lim J, Rose K-A, Anastasio K, Brower K, Glick L, Sharma L, Sewanan L, Knaggs L, Minasyan M, Batsu M, Petrone M, Kuang M, Nakahata M, Linehan M, Askenase MH, Simonov M, Smolgovsky M, Balkcom NC, Sonnert N, Naushad N, Vijayakumar P, Martinello R, Datta R, Handoko R, Bermejo S, Prophet S, Bickerton S, Velazquez S, Alpert T, Rice T, Khoury-Hanold W, Peng X, Yang Y, Cao Y, Strong Y, Lin Z, Wyllie AL, Campbell M, Lee AI, Chun HJ, Grubaugh ND, Schulz WL, Farhadian S, Dela Cruz C, Ring AM, Shaw AC, Wisnewski A V., Yildirim I, Ko AI, Omer SB, Iwasaki A. 2021. Delayed production of neutralizing antibodies correlates with fatal COVID-19. Nat Med 27:1178– 1186.

18. Garcia-Beltran WF, Lam EC, Astudillo MG, Yang D, Miller TE, Feldman J, Hauser BM, Caradonna TM, Clayton KL, Nitido AD, Murali MR, Alter G, Charles RC, Dighe A, Branda JA, Lennerz JK, Lingwood D, Schmidt AG, Iafrate AJ, Balazs AB. 2021. COVID-19-neutralizing antibodies predict disease severity and survival. Cell 184:476–488.e11.

19. MacLeod MKL, David A, McKee AS, Crawford F, Kappler JW, Marrack P. 2011. Memory CD4 T Cells That Express CXCR5 Provide Accelerated Help to B Cells. The Journal of Immunology 186:2889–2896.

20. Chen JS, Chow RD, Song E, Mao T, Israelow B, Kamath K, Bozekowski J, Haynes WA, Filler RB, Menasche BL, Wei J, Alfajaro MM, Song W, Peng L, Carter L, Weinstein JS, Gowthaman U, Chen S, Craft J, Shon JC, Iwasaki A, Wilen CB, Eisenbarth SC. 2022. High-affinity, neutralizing antibodies to SARS-CoV-2 can be made without T follicular helper cells. Sci Immunol 7:eabl5652.

21. Law JC, Koh WH, Budylowski P, Lin J, Yue F, Abe KT, Rathod B, Girard M, Li Z, Rini JM, Mubareka S, McGeer A, Chan AK, Gingras A-C, Watts TH, A. Ostrowski M. 2021. Systematic Examination of Antigen-Specific Recall T Cell Responses to SARS-CoV-2 versus Influenza Virus Reveals a Distinct Inflammatory Profile. The Journal of Immunology 206:37–50.

22. Law JC, Girard M, Chao GYC, Ward LA, Isho B, Rathod B, Colwill K, Li Z, Rini JM, Yue FY, Mubareka S, McGeer AJ, Ostrowski MA, Gommerman JL, Gingras A-C, Watts TH. 2022. Persistence of T Cell and Antibody Responses to SARS-CoV-2 Up to 9 Months after Symptom Onset. The Journal of Immunology 208:429–443.

23. Dayam RM, Law JC, Goetgebuer RL, Chao GYC, Abe KT, Sutton M, Finkelstein N, Stempak JM, Pereira D, Croitoru D, Acheampong L, Rizwan S, Rymaszewski K, Milgrom R, Ganatra D, Batista N V., Girard M, Lau I, Law R, Cheung MW, Rathod B, Kitaygorodsky J, Samson R, Hu Q, Hardy WR, Haroon N, Inman RD, Piguet V, Chandran V, Silverberg MS, Gingras A-C, Watts TH. 2022. Accelerated waning of immunity to SARS-CoV-2 mRNA vaccines in patients with immune-mediated inflammatory diseases. JCI Insight 7:e159721.

24. Cheung MW, Dayam RM, Shapiro JR, Law JC, Chao GYC, Pereira D, Goetgebuer RL, Croitoru D, Stempak JM, Acheampong L, Rizwan S, Lee JD, Jacob L, Ganatra D, Law R, Rodriguez-Castellanos VE, Kern-Smith M, Delgado-Brand M, Mailhot G, Haroon N, Inman RD, Piguet V, Chandran V, Silverberg MS, Watts TH, Gingras A-C. 2023. Third and Fourth Vaccine Doses Broaden and Prolong Immunity to SARS-CoV-2 in Adult Patients with Immune-Mediated Inflammatory Diseases. The Journal of Immunology 211:351–364.

25. Klingler Jã, Weiss S, Itri V, Liu X, Oguntuyo KY, Stevens C, Ikegame S, Hung CT, Enyindah-Asonye G, Amanat F, Baine I, Arinsburg S, Bandres JC, Kojic EM, Stoever J, Jurczyszak D, Bermudez-Gonzalez M, NÃidas A, Liu S, Lee B, Zolla-Pazner S, Hioe CE. 2021. Role of Immunoglobulin M and A Antibodies in the Neutralization of Severe Acute Respiratory Syndrome Coronavirus 2. Journal of Infectious Diseases 223:957–970.

26. Sheikh-Mohamed S, Isho B, Chao GYC, Zuo M, Cohen C, Lustig Y, Nahass GR, Salomon-Shulman RE, Blacker G, Fazel-Zarandi M, Rathod B, Colwill K, Jamal A, Li Z, de Launay KQ, Takaoka A, Garnham-Takaoka J, Patel A, Fahim C, Paterson A, Li AX, Haq N, Barati S, Gilbert L, Green K, Mozafarihashjin M, Samaan P, Budylowski P, Siqueira WL, Mubareka S, Ostrowski M, Rini JM, Rojas OL, Weissman IL, Tal MC, McGeer A, Regev-Yochay G, Straus S, Gingras AC, Gommerman JL. 2022. Systemic and mucosal IgA responses are variably induced in response to SARS-CoV-2 mRNA vaccination and are associated with protection against subsequent infection. Mucosal Immunol 15:799–808.

27. Mingari MC, Gerosa F, Carra G, Accolla RS, Moretta A, Zubler RH, Waldmann TA, Moretta L. 1984. Human interleukin-2 promotes proliferation of activated B cells via surface receptors similar to those of activated T cells. Nature 312:641–643.

28. Le Gallou S, Caron G, Delaloy C, Rossille D, Tarte K, Fest T. 2012. IL-2 Requirement for Human Plasma Cell Generation: Coupling Differentiation and Proliferation by Enhancing MAPK–ERK Signaling. The Journal of Immunology 189:161–173.

29. Gui L, Zeng Q, Xu Z, Zhang H, Qin S, Liu C, Xu C, Qian Z, Zhang S, Huang S, Chen L. 2016. IL-2, IL-4, IFN-γ or TNF-α enhances BAFF-stimulated cell viability and survival by activating Erk1/2 and S6K1 pathways in neoplastic B-lymphoid cells. Cytokine 84:37–46.

30. Miyauchi K, Sugimoto-Ishige A, Harada Y, Adachi Y, Usami Y, Kaji T, Inoue K, Hasegawa H, Watanabe T, Hijikata A, Fukuyama S, Maemura T, Okada-Hatakeyama M, Ohara O, Kawaoka Y, Takahashi Y, Takemori T, Kubo M. 2016. Protective neutralizing influenza antibody response in the absence of T follicular helper cells. Nat Immunol 17:1447–1458.

31. Lebman DA, Lee FD, Coffman RL. 1990. Mechanism for transforming growth factor beta and IL-2 enhancement of IgA expression in lipopolysaccharide-stimulated B cell cultures. The Journal of Immunology 144:952–959.

32. Spieker-Polet H, Yam P-C, Arbieva Z, Zhai S-K, Knight KL. 1999. In Vitro Induction of the Expression of Multiple IgA Isotype Genes in Rabbit B Cells by TGF-β and IL-2. The Journal of Immunology 162:5380–5388.

33. Murray SM, Ansari AM, Frater J, Klenerman P, Dunachie S, Barnes E, Ogbe A. 2023. The impact of pre-existing cross-reactive immunity on SARS-CoV-2 infection and vaccine responses. Nat Rev Immunol 23:304–316.

34. Mateus J, Grifoni A, Tarke A, Sidney J, Ramirez SI, Dan JM, Burger ZC, Rawlings SA, Smith DM, Phillips E, Mallal S, Lammers M, Rubiro P, Quiambao L, Sutherland A, Yu ED, Da Silva Antunes R, Greenbaum J, Frazier A, Markmann AJ, Premkumar L, De Silva A, Peters B, Crotty S, Sette A, Weiskopf D. 2020. Selective and cross-reactive SARS-CoV-2 T cell epitopes in unexposed humans. Science 370:89–94.

35. Braun J, Loyal L, Frentsch M, Wendisch D, Georg P, Kurth F, Hippenstiel S, Dingeldey M, Kruse B, Fauchere F, Baysal E, Mangold M, Henze L, Lauster R, Mall MA, Beyer K, Röhmel J, Voigt S, Schmitz J, Miltenyi S, Demuth I, Müller MA, Hocke A, Witzenrath M, Suttorp N, Kern F, Reimer U, Wenschuh H, Drosten C, Corman VM, Giesecke-Thiel C, Sander LE, Thiel A. 2020. SARS-CoV-2-reactive T cells in healthy donors and patients with COVID-19. Nature 587:270–274.

36. Peng Y, Mentzer AJ, Liu G, Yao X, Yin Z, Dong D, Dejnirattisai W, Rostron T, Supasa P, Liu C, López-Camacho C, Slon-Campos J, Zhao Y, Stuart DI, Paesen GC, Grimes JM, Antson AA, Bayfield OW, Hawkins DEDP, Ker DS, Wang B, Turtle L, Subramaniam K, Thomson P, Zhang P, Dold C, Ratcliff J, Simmonds P, de Silva T, Sopp P, Wellington D, Rajapaksa U, Chen YL, Salio M, Napolitani G, Paes W, Borrow P, Kessler BM, Fry JW, Schwabe NF, Semple MG, Baillie JK, Moore SC, Openshaw PJM, Ansari MA, Dunachie S, Barnes E, Frater J, Kerr G, Goulder P, Lockett T, Levin R, Zhang Y, Jing R, Ho LP, Dong T, Klenerman P, McMichael A, Ogg G, Kenneth Baillie J, Cornall RJ, Conlon CP, Screaton GR, Mongkolsapaya J, Knight JC. 2020. Broad and strong memory CD4+ and CD8+ T cells induced by SARS-CoV-2 in UK convalescent individuals following COVID-19. Nat Immunol 21:1336–1345.

37. Grifoni A, Weiskopf D, Ramirez SI, Mateus J, Dan JM, Moderbacher CR, Rawlings SA, Sutherland A, Premkumar L, Jadi RS, Marrama D, de Silva AM, Frazier A, Carlin AF, Greenbaum JA, Peters B, Krammer F, Smith DM, Crotty S, Sette A. 2020. Targets of T Cell Responses to SARS-CoV-2 Coronavirus in Humans with COVID-19 Disease and Unexposed Individuals. Cell 181:1489–1501.E15.

38. Ogbe A, Pace M, Bittaye M, Tipoe T, Adele S, Alagaratnam J, Aley PK, Ansari MA, Bara A, Broadhead S, Brown A, Brown H, Cappuccini F, Cinardo P, Dejnirattisai W, Ewer KJ, Fok H, Folegatti PM, Fowler J, Godfrey L, Goodman AL, Jackson B, Jenkin D, Jones M, Longet S, Makinson RA, Marchevsky NG, Mathew M, Mazzella A, Mujadidi YF, Parolini L, Petersen C, Plested E, Pollock KM, Rajeswaran T, Ramasamy MN, Rhead S, Robinson H, Robinson N, Sanders H, Serrano S, Tipton T, Waters A, Zacharopoulou P, Barnes E, Dunachie S, Goulder P, Klenerman P, Screaton GR, Winston A, Hill AVS, Gilbert SC, Carroll M, Pollard AJ, Fidler S, Fox J, Lambe T, Frater J. 2022. Durability of ChAdOx1 nCoV-19 vaccination in people living with HIV. JCI Insight 7:e157031.

39. Mateus J, Dan JM, Zhang Z, Moderbacher CR, Lammers M, Goodwin B, Sette A, Crotty S, Weiskopf D. 2021. Low-dose mRNA-1273 COVID-19 vaccine generates durable memory enhanced by cross-reactive T cells. Science 374:eabj9853.

40. Loyal L, Braun J, Henze L, Kruse B, Dingeldey M, Reimer U, Kern F, Schwarz T, Mangold M, Unger C, Dörfler F, Kadler S, Rosowski J, Gürcan K, Uyar-Aydin Z, Frentsch M, Kurth F, Schnatbaum K, Eckey M, Hippenstiel S, Hocke A, Müller MA, Sawitzki B, Miltenyi S, Paul F, Mall MA, Wenschuh H, Voigt S, Drosten C, Lauster R, Lachman N, Sander L-E, Corman VM, Röhmel J, Meyer-Arndt L, Thiel A, Giesecke-Thiel C. 2021. Cross-reactive CD4+ T cells enhance SARS-CoV-2 immune responses upon infection and vaccination. Science 374:eabh1823.

41. Berard M, Tough DF. 2002. Qualitative differences between naïve and memory T cells. Immunology 10.1046/j.1365-2567.2002.01447.x.

42. Ontario. 2021. Long Term Care Homes. https://tinyurl.com/nhe4exy7.

43. Government of Canada. 2021. COVID-19 daily epidemiology update. https://bit.ly/3kr7zQF.

44. Colwill K, Galipeau Y, Stuible M, Gervais C, Arnold C, Rathod B, Abe KT, Wang JH, Pasculescu A, Maltseva M, Rocheleau L, Pelchat M, Fazel-Zarandi M, Iskilova M, Barrios-Rodiles M, Bennett L, Yau K, Cholette F, Mesa C, Li AX, Paterson A, Hladunewich MA, Goodwin PJ, Wrana JL, Drews SJ, Mubareka S, McGeer AJ, Kim J, Langlois M, Gingras A, Durocher Y. 2022. A scalable serology solution for profiling humoral immune responses to SARS-CoV-2 infection and vaccination. Clin Transl Immunology 11:e1380.

45. Isho B, Abe KT, Zuo M, Jamal AJ, Rathod B, Wang JH, Li Z, Chao G, Rojas OL, Bang YM, Pu A, Christie-Holmes N, Gervais C, Ceccarelli D, Samavarchi-Tehrani P, Guvenc F, Budylowski P, Li A, Paterson A, Yue FY, Marin LM, Caldwell L, Wrana JL, Colwill K, Sicheri F, Mubareka S, Gray-Owen SD, Drews SJ, Siqueira WL, Barrios-Rodiles M, Ostrowski M, Rini JM, Durocher Y, McGeer AJ, Gommerman JL, Gingras A-C. 2020. Persistence of serum and saliva antibody responses to SARS-CoV-2 spike antigens in COVID-19 patients. Sci Immunol 5:eabe5511.

46. Amanat F, Stadlbauer D, Strohmeier S, Nguyen THO, Chromikova V, McMahon M, Jiang K, Arunkumar GA, Jurczyszak D, Polanco J, Bermudez-Gonzalez M, Kleiner G, Aydillo T, Miorin L, Fierer DS, Lugo LA, Kojic EM, Stoever J, Liu STH, Cunningham-Rundles C, Felgner PL, Moran T, García-Sastre A, Caplivski D, Cheng AC, Kedzierska K, Vapalahti O, Hepojoki JM, Simon V, Krammer F. 2020. A serological assay to detect SARS-CoV-2 seroconversion in humans. Nat Med 26:1033–1036.

47. Hollander M, Wolfe D, Chicken E. 2014. Nonparametric Statistical Methods 3rd Edition. John Wiley & Sons, Inc., Hoboken.

48. Ontario Agency for Health Protection and Promotion (Public Health Ontario). 2021. Enhanced epidemiological summary: COVID-19 variants of concern in Ontario: December 1, 2020 to May 9, 2021. Toronto.

49. Ontario Agency for Health Protection and Promotion (Public Health Ontario). 2022. Epidemiology summary: SARS-CoV-2 whole genome sequencing in Ontario, January 11, 2022. Toronto.

50. Hodcroft B E. 2024. CoVariants: SARS-CoV-2 Mutations and Variants of Interest. https://covariants.org/. Retrieved 25 April 2024.

51. Mveang Nzoghe A, Essone PN, Leboueny M, Maloupazoa Siawaya AC, Bongho EC, Mvoundza Ndjindji O, Avome Houechenou RM, Agnandji ST, Djoba Siawaya JF. 2021. Evidence and implications of pre-existing humoral cross-reactive immunity to SARS-CoV-2. Immun Inflamm Dis 9:128–133.

52. Anderson EM, Goodwin EC, Verma A, Arevalo CP, Bolton MJ, Weirick ME, Gouma S, McAllister CM, Christensen SR, Weaver JE, Hicks P, Manzoni TB, Oniyide O, Ramage H, Mathew D, Baxter AE, Oldridge DA, Greenplate AR, Wu JE, Alanio C, D’Andrea K, Kuthuru O, Dougherty J, Pattekar A, Kim J, Han N, Apostolidis SA, Huang AC, Vella LA, Kuri-Cervantes L, Pampena MB, Betts MR, Wherry EJ, Meyer NJ, Cherry S, Bates P, Rader DJ, Hensley SE. 2021. Seasonal human coronavirus antibodies are boosted upon SARS-CoV-2 infection but not associated with protection. Cell 184:1858–1864.e10.

53. Kundu R, Narean JS, Wang L, Fenn J, Pillay T, Fernandez ND, Conibear E, Koycheva A, Davies M, Tolosa-Wright M, Hakki S, Varro R, McDermott E, Hammett S, Cutajar J, Thwaites RS, Parker E, Rosadas C, McClure M, Tedder R, Taylor GP, Dunning J, Lalvani A. 2022. Cross-reactive memory T cells associate with protection against SARS-CoV-2 infection in COVID-19 contacts. Nat Commun 13:80.

54. Nelde A, Bilich T, Heitmann JS, Maringer Y, Salih HR, Roerden M, Lübke M, Bauer J, Rieth J, Wacker M, Peter A, Hörber S, Traenkle B, Kaiser PD, Rothbauer U, Becker M, Junker D, Krause G, Strengert M, Schneiderhan-Marra N, Templin MF, Joos TO, Kowalewski DJ, Stos-Zweifel V, Fehr M, Rabsteyn A, Mirakaj V, Karbach J, Jäger E, Graf M, Gruber LC, Rachfalski D, Preuß B, Hagelstein I, Märklin M, Bakchoul T, Gouttefangeas C, Kohlbacher O, Klein R, Stevanović S, Rammensee HG, Walz JS. 2021. SARS-CoV-2-derived peptides define heterologous and COVID-19-induced T cell recognition. Nat Immunol 22:74–85.

55. Isho B, Abe KT, Zuo M, Jamal AJ, Rathod B, Wang JH, Li Z, Chao G, Rojas OL, Bang YM, Pu A, Christie-Holmes N, Gervais C, Ceccarelli D, Samavarchi-Tehrani P, Guvenc F, Budylowski P, Li A, Paterson A, Yue FY, Marin LM, Caldwell L, Wrana JL, Colwill K, Sicheri F, Mubareka S, Gray-Owen SD, Drews SJ, Siqueira WL, Barrios-Rodiles M, Ostrowski M, Rini JM, Durocher Y, McGeer AJ, Gommerman JL, Gingras A-C. 2020. Persistence of serum and saliva antibody responses to SARS-CoV-2 spike antigens in COVID-19 patients. Sci Immunol 5:eabe5511.

56. Dan JM, Mateus J, Kato Y, Hastie KM, Yu ED, Faliti CE, Grifoni A, Ramirez SI, Haupt S, Frazier A, Nakao C, Rayaprolu V, Rawlings SA, Peters B, Krammer F, Simon V, Saphire EO, Smith DM, Weiskopf D, Sette A, Crotty S. 2021. Immunological memory to SARS-CoV-2 assessed for up to 8 months after infection. Science 371:eabf4063.

57. Wisnewski A V., Campillo Luna J, Redlich CA. 2021. Human IgG and IgA responses to COVID-19 mRNA vaccines. PLoS One 16:e0249499.

58. Hurme A, Jalkanen P, Heroum J, Liedes O, Vara S, Melin M, Teräsjärvi J, He Q, Pöysti S, Hänninen A, Oksi J, Vuorinen T, Kantele A, Tähtinen PA, Ivaska L, Kakkola L, Lempainen J, Julkunen I. 2022. Long-Lasting T Cell Responses in BNT162b2 COVID-19 mRNA Vaccinees and COVID-19 Convalescent Patients. Front Immunol 13:869990.

59. Maringer Y, Nelde A, Schroeder SM, Schuhmacher J, Hörber S, Peter A, Karbach J, Jäger E, Walz JS. 2022. Durable spike-specific T cell responses after different COVID-19 vaccination regimens are not further enhanced by booster vaccination. Sci Immunol 7:eadd3899.

60. Burns MD, Boribong BP, Bartsch YC, Loiselle M, St. Denis KJ, Sheehan ML, Chen JW, Davis JP, Lima R, Edlow AG, Fasano A, Balazs AB, Alter G, Yonker LM. 2022. Durability and Cross-Reactivity of SARS-CoV-2 mRNA Vaccine in Adolescent Children. Vaccines (Basel) 10:492.

61. Cheung MW, Dayam RM, Shapiro JR, Law JC, Chao GYC, Pereira D, Goetgebuer RL, Croitoru D, Stempak JM, Acheampong L, Rizwan S, Lee JD, Jacob L, Ganatra D, Law R, Rodriguez-Castellanos VE, Kern-Smith M, Delgado-Brand M, Mailhot G, Haroon N, Inman RD, Piguet V, Chandran V, Silverberg MS, Watts TH, Gingras A-C. 2023. Third and Fourth Vaccine Doses Broaden and Prolong Immunity to SARS-CoV-2 in Adult Patients with Immune-Mediated Inflammatory Diseases. The Journal of Immunology 211:351–364.

62. Matveev VA, Mihelic EZ, Benko E, Budylowski P, Grocott S, Lee T, Korosec CS, Colwill K, Stephenson H, Law R, Ward LA, Sheikh-Mohamed S, Mailhot G, Delgado-Brand M, Pasculescu A, Wang JH, Qi F, Tursun T, Kardava L, Chau S, Samaan P, Imran A, Copertino DC, Chao G, Choi Y, Reinhard RJ, Kaul R, Heffernan JM, Jones RB, Chun T-W, Moir S, Singer J, Gommerman J, Gingras A-C, Kovacs C, Ostrowski M. 2023. Immunogenicity of COVID-19 vaccines and their effect on HIV reservoir in older people with HIV. iScience 26:107915.

63. Goel RR, Painter MM, Apostolidis SA, Mathew D, Meng W, Rosenfeld AM, Lundgreen KA, Reynaldi A, Khoury DS, Pattekar A, Gouma S, Kuri-Cervantes L, Hicks P, Dysinger S, Hicks A, Sharma H, Herring S, Korte S, Baxter AE, Oldridge DA, Giles JR, Weirick ME, McAllister CM, Awofolaju M, Tanenbaum N, Drapeau EM, Dougherty J, Long S, D’Andrea K, Hamilton JT, McLaughlin M, Williams JC, Adamski S, Kuthuru O, Frank I, Betts MR, Vella LA, Grifoni A, Weiskopf D, Sette A, Hensley SE, Davenport MP, Bates P, Luning Prak ET, Greenplate AR, Wherry EJ. 2021. mRNA vaccines induce durable immune memory to SARS-CoV-2 and variants of concern. Science 374:abm0829.

64. Naaber P, Tserel L, Kangro K, Sepp E, Jürjenson V, Adamson A, Haljasmägi L, Rumm AP, Maruste R, Kärner J, Gerhold JM, Planken A, Ustav M, Kisand K, Peterson P. 2021. Dynamics of antibody response to BNT162b2 vaccine after six months: a longitudinal prospective study. The Lancet Regional Health - Europe 10:100208.

65. Dayam RM, Law JC, Goetgebuer RL, Chao GYC, Abe KT, Sutton M, Finkelstein N, Stempak JM, Pereira D, Croitoru D, Acheampong L, Rizwan S, Rymaszewski K, Milgrom R, Ganatra D, Batista N V., Girard M, Lau I, Law R, Cheung MW, Rathod B, Kitaygorodsky J, Samson R, Hu Q, Hardy WR, Haroon N, Inman RD, Piguet V, Chandran V, Silverberg MS, Gingras A-C, Watts TH. 2022. Accelerated waning of immunity to SARS-CoV-2 mRNA vaccines in patients with immune-mediated inflammatory diseases. JCI Insight 7:e159721.

66. Swadling L, Diniz MO, Schmidt NM, Amin OE, Chandran A, Shaw E, Pade C, Gibbons JM, Le Bert N, Tan AT, Jeffery-Smith A, Tan CCS, Tham CYL, Kucykowicz S, Aidoo-Micah G, Rosenheim J, Davies J, Johnson M, Jensen MP, Joy G, McCoy LE, Valdes AM, Chain BM, Goldblatt D, Altmann DM, Boyton RJ, Manisty C, Treibel TA, Moon JC, Abbass H, Abiodun A, Alfarih M, Alldis Z, Andiapen M, Artico J, Augusto JB, Baca GL, Bailey SNL, Bhuva AN, Boulter A, Bowles R, Boyton RJ, Bracken O V., O’Brien B, Brooks T, Bullock N, Butler DK, Captur G, Champion N, Chan C, Collier D, de Sousa JC, Couto-Parada X, Cutino-Mogue T, Davies RH, Douglas B, Di Genova C, Dieobi-Anene K, Ellis A, Feehan K, Finlay M, Fontana M, Forooghi N, Gaier C, Gilroy D, Hamblin M, Harker G, Hewson J, Hickling LM, Hingorani AD, Howes L, Hughes A, Hughes G, Hughes R, Itua I, Jardim V, Lee W-YJ, Jensen MP, Jones J, Jones M, Joy G, Kapil V, Kurdi H, Lambourne J, Lin K-M, Louth S, Mandadapu V, McKnight Á, Menacho K, Mfuko C, Mitchelmore O, Moon C, Munoz-Sandoval D, Murray SM, Noursadeghi M, Otter A, Palma S, Parker R, Patel K, Pawarova B, Petersen SE, Piniera B, Pieper FP, Pope D, Prossora M, Rannigan L, Rapala A, Reynolds CJ, Richards A, Robathan M, Sambile G, Semper A, Seraphim A, Simion M, Smit A, Sugimoto M, Taylor S, Temperton N, Thomas S, Thornton GD, Tucker A, Veerapen J, Vijayakumar M, Welch S, Wodehouse T, Wynne L, Zahedi D, van Dorp L, Balloux F, McKnight Á, Noursadeghi M, Bertoletti A, Maini MK. 2022. Pre-existing polymerase-specific T cells expand in abortive seronegative SARS-CoV-2. Nature 601:110–117.

67. Wang Z, Yang X, Zhong J, Zhou Y, Tang Z, Zhou H, He J, Mei X, Tang Y, Lin B, Chen Z, McCluskey J, Yang J, Corbett AJ, Ran P. 2021. Exposure to SARS-CoV-2 generates T-cell memory in the absence of a detectable viral infection. Nat Commun 12:1724.

68. Moss P. 2022. The T cell immune response against SARS-CoV-2. Nat Immunol 23:186–193.

69. Cheung MW, Dayam RM, Shapiro JR, Law JC, Chao GYC, Pereira D, Goetgebuer RL, Croitoru D, Stempak JM, Acheampong L, Rizwan S, Lee JD, Jacob L, Ganatra D, Law R, Rodriguez-Castellanos VE, Kern-Smith M, Delgado-Brand M, Mailhot G, Haroon N, Inman RD, Piguet V, Chandran V, Silverberg MS, Watts TH, Gingras A-C. 2023. Third and Fourth Vaccine Doses Broaden and Prolong Immunity to SARS-CoV-2 in Adult Patients with Immune-Mediated Inflammatory Diseases. The Journal of Immunology 211:351–364.

70. Chen Y, Tong P, Whiteman N, Sanjari Moghaddam A, Zarghami M, Zuiani A, Habibi S, Gautam A, Keerti, Bi C, Xiao T, Cai Y, Chen B, Neuberg D, Wesemann DR. 2022. Immune recall improves antibody durability and breadth to SARS-CoV-2 variants. Sci Immunol 7:eabp8328.

71. Siracusano G, Ruggiero A, Bisoffi Z, Piubelli C, Carbonare LD, Valenti MT, Mayora-Neto M, Temperton N, Lopalco L, Zipeto D. 2022. Different decay of antibody response and VOC sensitivity in naïve and previously infected subjects at 15 weeks following vaccination with BNT162b2. J Transl Med 20:22.

72. Buckner CM, Kardava L, El Merhebi O, Narpala SR, Serebryannyy L, Lin BC, Wang W, Zhang X, Lopes de Assis F, Kelly SEM, Teng I-T, McCormack GE, Praiss LH, Seamon CA, Rai MA, Kalish H, Kwong PD, Proschan MA, McDermott AB, Fauci AS, Chun T-W, Moir S. 2022. Interval between prior SARS-CoV-2 infection and booster vaccination impacts magnitude and quality of antibody and B cell responses. Cell 185:4333–4346.e14.

73. Brunet-Ratnasingham E, Morin S, Randolph HE, Labrecque M, Bélair J, Lima-Barbosa R, Pagliuzza A, Marchitto L, Hultström M, Niessl J, Cloutier R, Sreng Flores AM, Brassard N, Benlarbi M, Prévost J, Ding S, Anand SP, Sannier G, Marks A, Wågsäter D, Bareke E, Zeberg H, Lipcsey M, Frithiof R, Larsson A, Zhou S, Nakanishi T, Morrison D, Vezina D, Bourassa C, Gendron-Lepage G, Medjahed H, Point F, Richard J, Larochelle C, Prat A, Cunningham JL, Arbour N, Durand M, Richards JB, Moon K, Chomont N, Finzi A, Tétreault M, Barreiro L, Wolf G, Kaufmann DE. 2024. Sustained IFN signaling is associated with delayed development of SARS-CoV-2-specific immunity. Nat Commun 15:4177.

74. Radbruch A, McGrath MA, Siracusa F, Hoffmann U, Sercan-Alp Ö, Hutloff A, Tokoyoda K, Chang H-D, Dong J. 2021. Homeostasis and Durability of T-Cell Memory—The Resting and the Restless T-Cell Memory. Cold Spring Harb Perspect Biol 13:a038083.

75. Tokoyoda K, Zehentmeier S, Hegazy AN, Albrecht I, Grün JR, Löhning M, Radbruch A. 2009. Professional Memory CD4+ T Lymphocytes Preferentially Reside and Rest in the Bone Marrow. Immunity 30:721–730.

76. Pacheco Y, Acosta-Ampudia Y, Monsalve DM, Chang C, Gershwin ME, Anaya J-M. 2019. Bystander activation and autoimmunity. J Autoimmun 103:102301.

77. Doisne J-M, Urrutia A, Lacabaratz-Porret C, Goujard C, Meyer L, Chaix M-L, Sinet M, Venet A. 2004. CD8+ T Cells Specific for EBV, Cytomegalovirus, and Influenza Virus Are Activated during Primary HIV Infection. The Journal of Immunology 173:2410–2418.

78. van Aalst S, Ludwig IS, van der Zee R, van Eden W, Broere F. 2017. Bystander activation of irrelevant CD4+ T cells following antigen-specific vaccination occurs in the presence and absence of adjuvant. PLoS One 12:e0177365.

